# Thermal Nano-Engineering of Ginger Extracellular Vesicles for Targeted Oral Therapy of Colitis

**DOI:** 10.64898/2026.02.24.707840

**Authors:** Linhai Hou, Jie Cao, Shengjie Gao, Xiaofan Wang, Zhongxian Zhang, Meiqi Li, Yu Mao, Changhong Liu, Ling Yan, Haiping Hao, Lei Zheng

## Abstract

Plant-derived extracellular vesicles (PEVs) are promising candidates for oral drug delivery, yet their clinical translation is hindered by limited targeting precision and inconsistent systemic absorption. While surface engineering can enhance tissue accumulation, strategies that preserve biocompatibility and enable scalable production remain limited. Here, we introduce a simple thermal processing approach, boiling, to structurally reconfigure ginger EVs into functionally enhanced, thermally reassembled nanoparticles (B-GEVs). The surface architecture of B-GEVs is enriched with key vesicle trafficking regulators, including V-type proton ATPase subunit G, ARF1, and β-adaptin-like protein. This specific composition drives their tissue-specific accumulation in the intestine and liver and potentiates clathrin-dependent cellular uptake in intestinal cells by 8.57-fold. Beyond superior intrinsic anti-inflammatory activity through NLRP3 inflammasome suppression, B-GEVs function as an efficient oral delivery platform. When loaded with TNF-α siRNA, they enable a synergistic therapy that simultaneously modulates upstream inflammation and silences key downstream mediators, showing potent efficacy in colitis. Our findings position boiling as a natural strategy for enhancing the bioactivity and targeted oral delivery potential of ginger-derived EVs.

## Introduction

Extracellular vesicles (EVs) are key mediators of intercellular communication and have gained increasing interest as potential therapeutic vehicles (*1*). Nature provides a rich source of inspiration for optimizing their delivery, including EVs derived from edible plants, which have evolved to facilitate nutrient assimilation (*2*). Over the course of human history, various vegetables have been consumed both in raw and cooked forms. Early investigations indicated that cooking solely alters phytochemical content and physical properties (*3*). Notably, vegetable juice represents a complex multicomponent system characterized by inherently heterogeneous structural assemblies rather than discrete single compounds (*4*). Among these, plant-derived EVs are key mediators of the bioactivity of vegetable juices (*5*); however, the consequences of boiling on the nanoscale structural and function of PEVs remain poorly elucidated. Thermal processing of plant matrices is known to elicit profound biochemical transformations (*6*, *7*). Yet, whether similar structural remodeling occurs within the architectures of GEVs during such processing remains an open question.

Ginger (*Zingiber officinale*) holds a prominent place in global culinary and medicinal traditions, consumed extensively both raw and cooked for over 5000 years (*8*). A growing body of clinical and preclinical studies has investigated the therapeutic potential of ginger supplementation, ginger-derived extracts, and ginger-derived extracellular vesicles (GEVs) in the management of diseases such as inflammatory bowel disease (IBD) (*5*, *9*, *10*, *11*). Prior research contrasting raw versus processed ginger has revealed substantial differences in the composition and bio-efficacy of its bioactive molecules (*11*, *12*). GEVs mediate intercellular communication through cargo transfer or membrane contact.

The therapeutic and delivery efficacy of PEVs, including GEVs, depend critically on their cellular uptake dynamics by delivering functional cargo, which vary based on EVs subtype and recipient cells (*9*, *13*, *14*). The surface of EVs incorporates proteins and lipids derived from their parental cells. Following isolation, EVs can subsequently acquire a dynamic protein corona from the extracellular environment, which collectively shapes their biological function, cellular uptake, and in vivo distribution (*15*, *16*). Therapeutic hypothermia induces temperature-dependent enrichment of apolipoproteins C1 in the protein corona of the nanocarrier surface, facilitating receptor-mediated targeting capability (*17*). Given that biological systems frequently exhibit stimulus-responsive structural reorganization in aqueous environments (*18*, *19*), we hypothesized that boiling—a common thermal process—could drive structural reorganization of GEVs, potentially enhancing their functional performance for targeted delivery.

In this study, we discovered that boiling triggers the nanoscale reconstitution of GEVs within ginger juice, forming a protein-rich shell that incorporates bioactive components and metal ions from the ginger matrix. This thermally driven structural remodeling substantially enhances GEVs internalization by intestinal epithelial and hepatic cells through clathrin-mediated endocytosis, while concurrently altering their *in vivo* biodistribution profiles. Notably, the reconfigured GEVs exhibited significantly stronger anti-inflammatory efficacy compared to native vesicles. Beyond their standalone activity, B-GEVs serve as a highly efficient oral-delivery platform. By delivering TNF-α siRNA, they facilitate a synergistic therapy that simultaneously targets upstream inflammatory pathways and silences key downstream mediators. Overall, boiling may serve as a natural nano-structural engineering approach, enhancing the functionality of plant-derived EVs and improving their potential for oral gene delivery and therapeutic efficacy.

## Results

### Liver and colon show preference for B-GEVs *in vivo*

EVs represent critical functional components of vegetable juices, acting as pivotal mediators of interspecies communication through the transfer of bioactive molecules across biological systems (*4*, *10*, *20*). To elucidate the effects of boiling processing on the biodistribution patterns of GEVs, we isolated EVs from fresh (F-GEVs) and boiled (B-GEVs) ginger juice (**Fig. 1a**). Transmission electron microscopy (TEM) revealed both F-GEVs and B-GEVs to display characteristic spherical morphologies consistent with previously described GEVs features (*9*, *21*) (**Fig. 1b**). Nanoparticle tracking analysis (NTA) demonstrated a 14 nm increase in average particle diameter in B-GEVs relative to F-GEVs (**Fig. 1c**), indicating heat-induced surface modifications. Given the selective organotropism of PEVs and their therapeutic potential (*21*, *22*, *23*), we evaluated the effects of boiling on *in vivo* GEVs biodistribution using DiR-labeled vesicles administered orally to mice. Quantitative imaging revealed that B-GEVs exhibited markedly enhanced hepatic and colonic accumulation, with 2.0-fold and 2.6-fold greater localization in the liver and colon, respectively, at 18 h post-administration, compared to F-GEVs (**Fig. 1d-g**). To further visualize the tissue distribution of F-GEVs and B-GEVs, fluorescence imaging was performed on organs isolated following oral administration. We found that B-GEVs accumulated more significantly in the intestine and liver compared to F-GEVs (**Fig. 1h and i**).

**Fig. 1.**
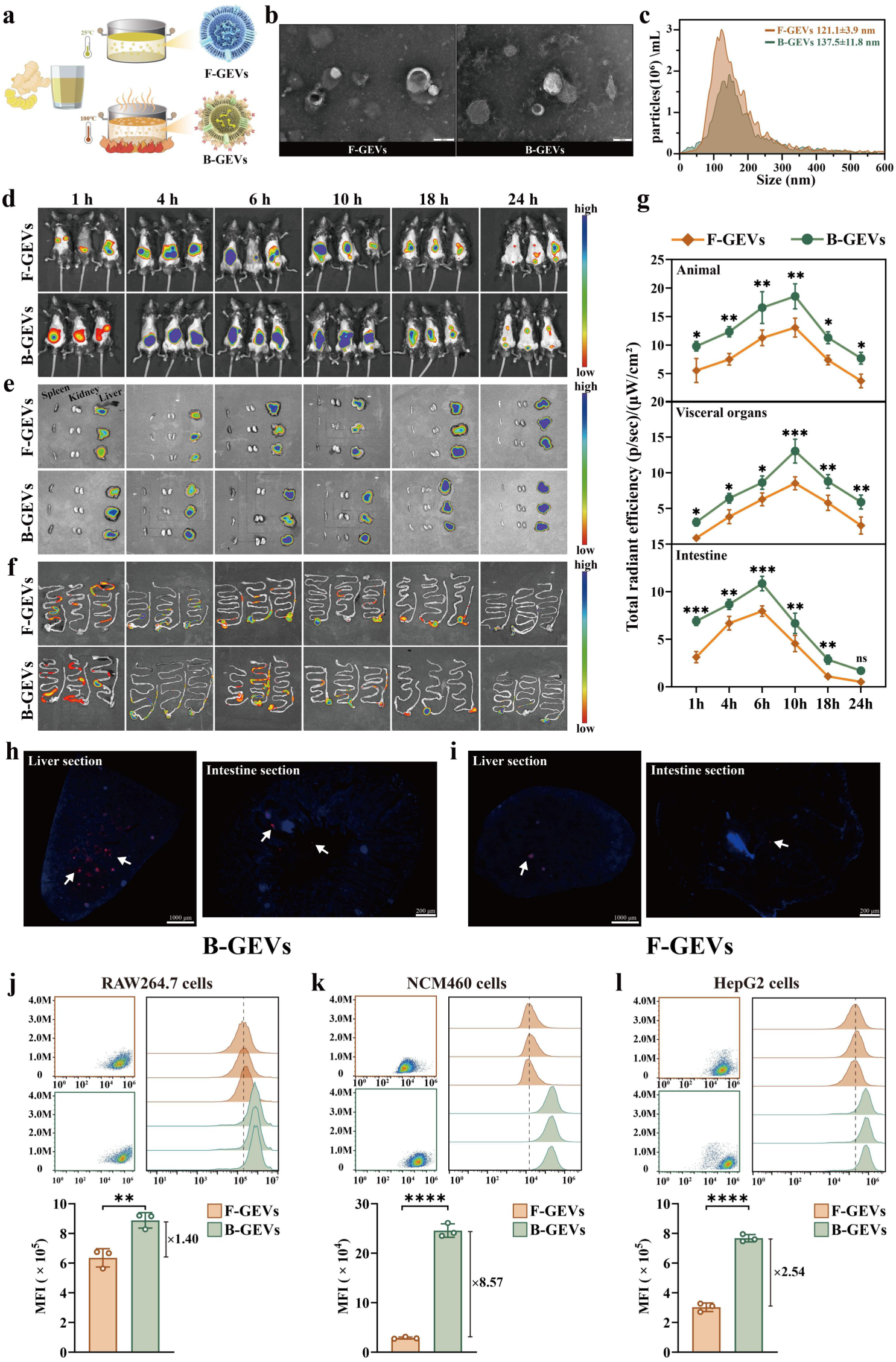
Targeted distribution and cellular uptake of B-GEVs. (**a)**, Schematic representation of the thermally processed GEVs. (**b)**, TEM images of F-GEVs and B-GEVs. **c**, Size distribution analysis of F-GEVs and B-GEVs. n=3. (**d)**, Fluorescence intensity distribution in mice following oral administration of F-GEVs and B-GEVs. (**e and f)**, Ex vivo organ imaging (spleen, kidney, liver, and intestine) at various time points post-administration. **(g)**, Quantification of liver and colon fluorescence signals (mean ± SD, two-tailed t-test). n=3 biologically independent samples. * *P* < 0.05, ** *P* < 0.01, *** *P* < 0.001. **(h and i)**, Confocal microscopic imaging of intestine and liver sections following oral administration of F-GEVs and B-GEVs. **(j-l)**, Flow cytometric analysis of the F-GEVs and B-GEVs cellular internalization by RAW 264.7 cells (**j**), NCM460 cells (**k**), and HepG2 cells (**l)**. n=3 biological replicates. Data are presented as means ± SD, n = 3. P values were calculated using two-tailed t-tests. ** *P* < 0.01, **** *P* < 0.0001, ns, non-significant.

To assess whether enhanced organ targeting correlated with improved cellular uptake, we quantified B-GEVs internalization in macrophages, hepatocytes, and intestinal epithelial cells. Viability assays confirmed no cytotoxicity at 5×10⁹ particles/mL for either B-GEVs or F-GEVs (**Fig. S1**). Uptake analyses revealed that B-GEVs exhibited 8.57-fold, 2.54-fold, and 1.40-fold higher internalization in NCM460 intestinal epithelial cells, HepG2 hepatocytes, and RAW 264.7 macrophages, respectively, relative to F-GEVs (**Fig. 1j-l and Fig. S2**). Confocal microscopy further confirmed the superior internalization of B-GEVs in RAW264.7 cells (**Fig. S3**). Collectively, the results suggested that B-GEVs exhibit favorable uptake profiles within the liver and intestinal microenvironment, as evidenced by increased internalization by phagocytes, hepatocytes, and intestinal epithelial cells.

### Boiling processing induces structural and compositional changes in GEVs

Biological systems assemble organic-inorganic complexes in response to thermal stimuli (*24*). Atomic force microscopy (AFM) revealed that B-GEVs displayed markedly increased surface roughness and heterogeneity compared to F-GEVs (**Fig. 2a**). Particle counts showed significantly more F-GEVs (1.3×10^11^ particles/mL) than B-GEVs (7.4×10^10^ particles/mL) (**Fig. S4**). Notably, protein content measurements demonstrated a 41% elevation in B-GEVs (2336.63 ± 230.77 ng/μL) relative to F-GEVs (1660.17 ± 193.93 ng/μL; **Fig. 2b**). Inductively coupled plasma mass spectrometry (ICP-MS) analysis revealed significantly increased levels of calcium (1.54-fold), manganese (1.24-fold), magnesium (1.49-fold), copper (1.61-fold), zinc (1.54-fold), and potassium (1.65-fold) in B-GEVs compared to F-GEVs (**Fig. 2d**). Boiling also led to a reduction in miRNA content within GEVs relative to the raw preparation **(Fig. S5)**.

**Fig. 2.**
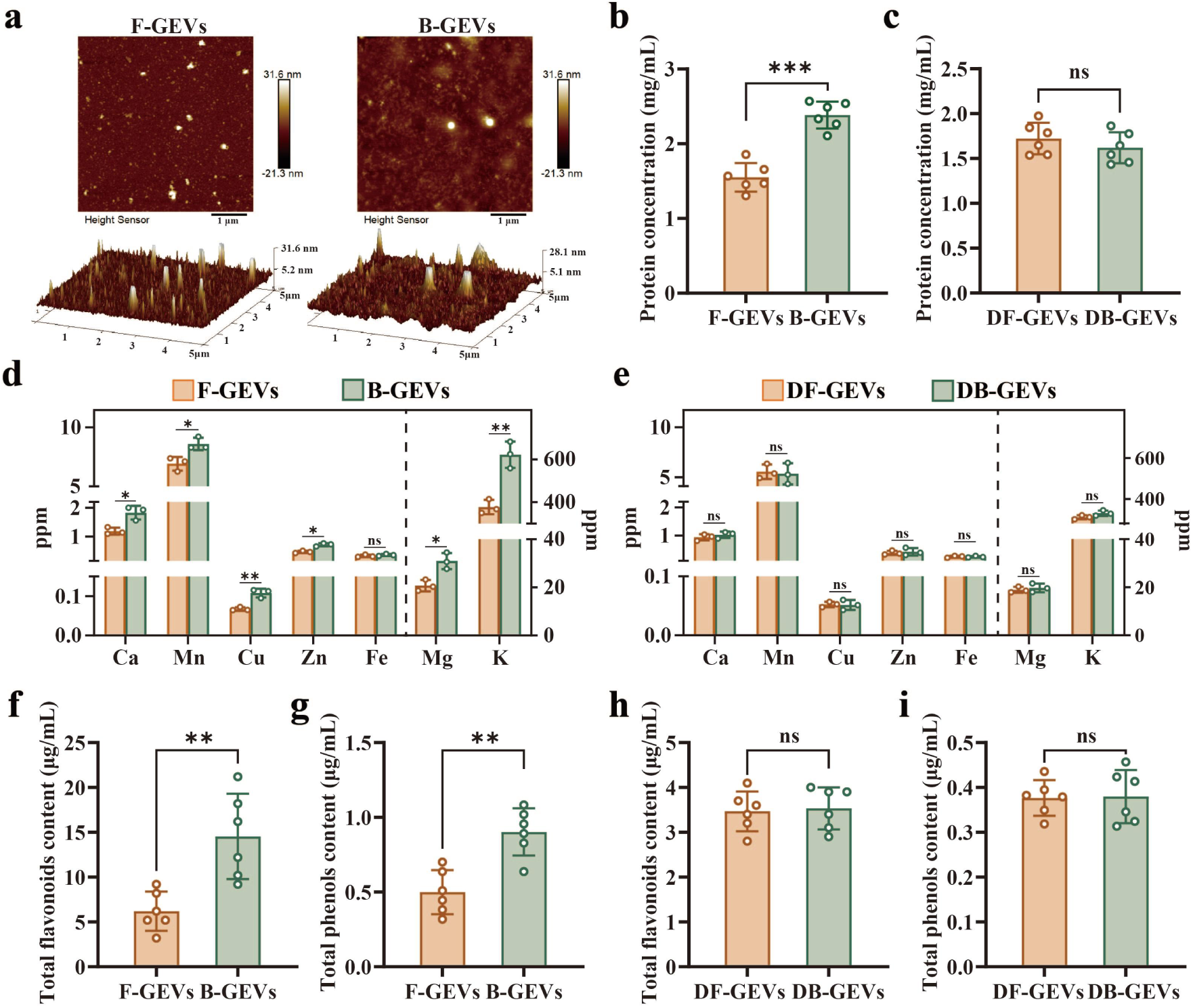
Boiling processing alters the structure and composition of GEVs. **(a)**, AFM topography maps of F-GEVs and B-GEVs. (**b and c)**, Protein quantification of F-GEVs and B-GEVs (b), as well as their digested forms DF-GEVs and DB-GEVs (c) by BCA assay. n=3 replicates. The two-sided unpaired t-test method was used for statistical significance testing. *** *P* < 0.001, ns, non-significant. (**d and e)**, ICP-MS analysis of physiological metal elements (Ca, Mn, Cu, Zn, Fe, Mg, and K) in both F-GEVs and B-GEVs (d), as well as DF-GEVs and DB-GEVs (e). n=3 replicates. The two-sided unpaired t-test method was used for statistical significance testing. * *P* < 0.05, ** *P* < 0.01, ns, non-significant. (**f-i)**, The measurement of total flavonoids and total phenols content in both F-GEVs and B-GEVs (f, g), as well as DF-GEVs and DB-GEVs (h, i) by UV-Vis spectrophotometry. n=3 replicates. The two-sided unpaired t-test method was used for statistical significance testing. ** *P* < 0.01, ns, non-significant.

To assess the origin of surface-bound proteins and ions, protease digestion of surface proteins on both F-GEVs and B-GEVs to yield DF-GEVs and DB-GEVs. Strikingly, protein and metal ion levels in DB-GEVs returned to those observed in DF-GEVs (**Fig. 2c-e**). The results imply that the increased metal ions on B-GEVs may be mainly bound to groups on ginger proteins via chelation or self-assembly, which can be transferred intercellularly to modulate biological responses (*25*). Furthermore, B-GEVs retained 1.8-fold higher total phenolic and flavonoid content compared to F-GEVs (**Fig. 2f and g**), a phenomenon absent in DB-GEVs (**Fig. 2h and i**). These findings indicate that boiling drives protein-phytochemical co-assembly through enhanced hydrophobic interactions, creating stabilized metal complexes that boost bioactive retention.

### Protein Shell formed on the surface of EVs isolated from boiling processing-treated ginger juice

To determine whether boiling induces protein adsorption onto GEVs, we compared surface-deproteinized samples of B-GEVs and F-GEVs, which exhibited comparable levels of residual protein (**Fig. 2c**). This finding indicates that the higher protein content observed in B-GEVs stems from surface-adsorbed proteins, not from intrinsic compositional differences. Time-course analysis demonstrated that extended boiling progressively increased both protein content and mechanical rigidity of GEVs (**Fig. 3a and b**), with B-GEVs exhibiting a 1.85-fold higher Young’s modulus after 60 min of thermal treatment (7.86 ± 1.04 MPa) compared to F-GEVs (4.25 ± 1.92 MPa) (**Fig. 3b**). Fluorescence quantification confirmed time-dependent accumulation of surface-adsorbed proteins (**Fig. 3c and d**). Direct visualization using fluorescently labeled proteins incubated with GEVs under thermal conditions revealed the formation of a distinct protein shell on B-GEVs surfaces (**Fig. 3e**). Thermal stress induced more changes in the secondary structure of proteins in solution. Circular dichroism (CD) spectra of ginger protein at various temperatures indicate that the protein is also primarily composed of α-helices (208 and 222 nm) and that the secondary structure is gradually disrupted as the temperature is increased (**Fig. 3f**). Upon heating, the β-sheet ratio of ginger protein was the lower. Furthermore, spectral changes in the 190–200 nm region upon boiling suggested the formation of non-native, partially ordered intermediates or aggregates, rather than a complete transition to a random-coil conformation (**Fig. 3f**). To further investigate the interaction between structural changes of ginger protein and GEVs, the thermodynamic response of protein adsorption on GEVs surfaces was determined using isothermal titration calorimetry (ITC). Titration thermograms showed a markedly stronger exothermic response for boiled proteins than for their native counterparts (**Fig. 3g**). The cumulative interaction heat (ΣΔQ₁–₁₉) increased from 207.7 μJ to 225.1 μJ after thermal treatment, indicating a more favorable enthalpic drive for assembly (**Fig. 3h**). Collectively, these results demonstrate that thermally unfolded proteins promote energetically favorable associations with the F-GEVs lipid bilayer, culminating in the formation of a stabilized protein shell on GEVs (**Fig. 3i**).

**Fig. 3.**
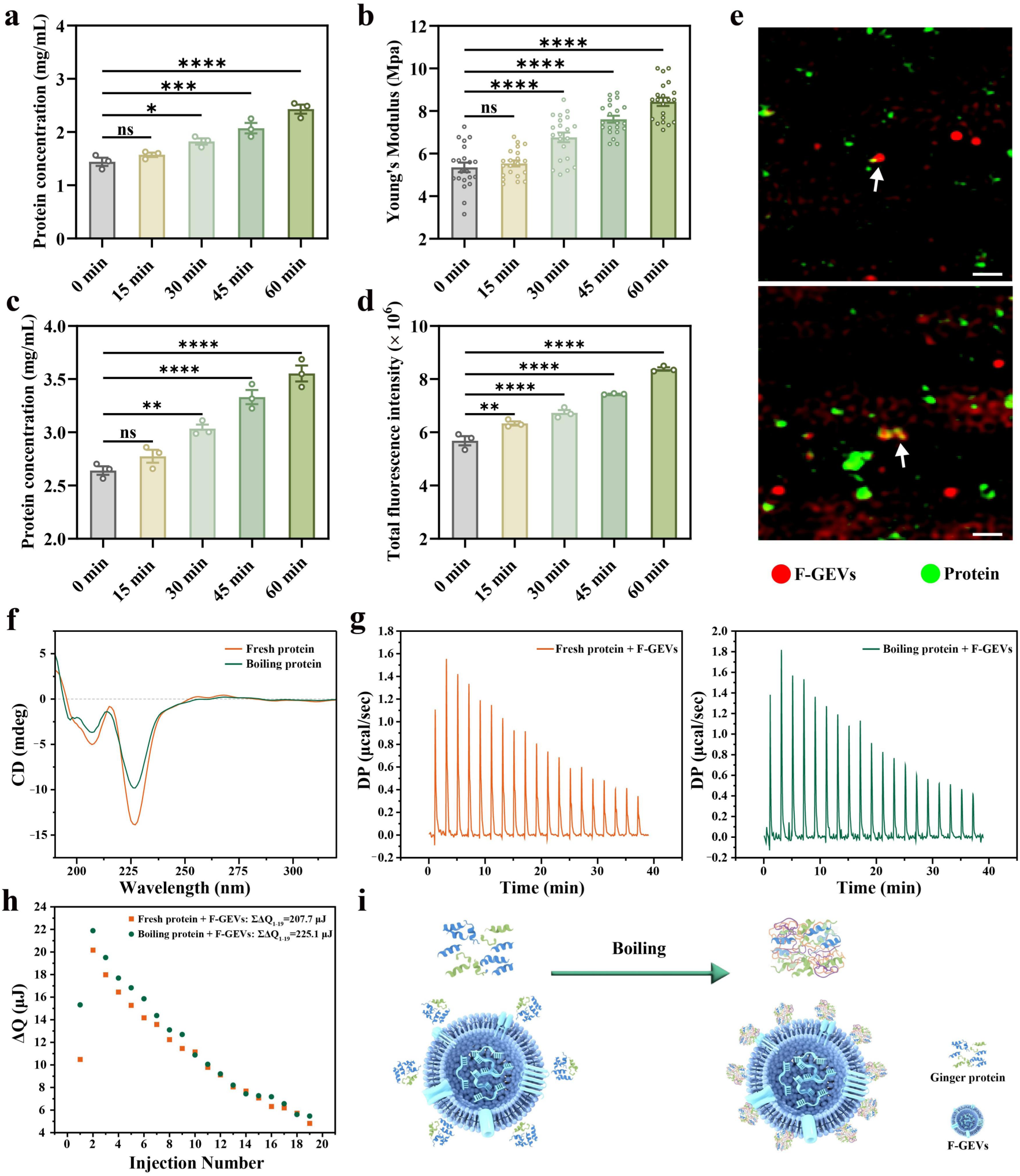
The protein shell formed on the surface of B-GEVs. **(a)**, Time-dependent protein adsorption of B-GEVs during boiling processing (0-60 min). **(b)**, AFM-measured Young’s modulus (n=20 particles per group). **(c)**, Protein concentrations after incubation of F-GEVs with fluorescently labeled proteins under controlled heating. **(d)**, Total fluorescence value of B-GEVs at various thermal treatment time points. **(e),** Confocal microscopy of Alex Fluor 488-labeled protein (green) co-localizing with DiD-labeled GEVs (red). Scale bars: 500 nm. **(f),** CD spectra of fresh and boiled ginger proteins. **(g),** ITC thermograms of fresh (left) and boiled (right) ginger proteins titrated into an F-GEVs suspension. **(h),** cumulative interaction heat. **(i),** Schematic illustration of ginger protein organization around F-GEVs before and after boiling. All data are presented as means ± SD, n = 3. P values were calculated using two-sided one-way ANOVA post-Dunnett’s test; \*\**P* < 0.01, \*\*\**P* < 0.001, \*\*\*\**P* < 0.0001, ns, non-significant.

### Boiling-induced remodeling of the GEVs proteome

To investigate boiling processing-driven proteomic remodeling in GEVs, quantitative proteomics via LC-MS/MS revealed substantial differences in protein composition between F-GEVs and B-GEVs (**Fig. 4**). SDS–PAGE demonstrated altered protein abundance in B-GEVs (**Fig. 4a**), with molecular weight distributions predominantly spanning 0-50 kDa (**Fig. 4b**). Lower-molecular-weight proteins, owing to higher diffusion coefficients, may preferentially bind B-GEVs surfaces. LC-MS/MS results showed minimal mass error and strong correlation among replicates, confirming data reliability (**Fig. S6a and b**). In total, 2706 proteins were identified in F-GEVs and 1010 proteins in B-GEVs, with no unique protein signatures exclusive to B-GEVs (**Fig. S7**), suggesting reorganization rather than acquisition of novel proteins. Volcano plot and clustering analyses identified 163 proteins significantly upregulated in B-GEVs (**Fig. 4c and d**), including key regulators of vesicle transport such as V-type proton ATPase subunit G, ADP-ribosylation factor 1 (ARF1), and β-adaptin-like protein (**Fig. 4e**). Additional proteins included those involved in immunity, metabolism, signaling, chaperone function, and protein storage (**Table S1**). Gene Ontology (GO) and Kyoto Encyclopedia of Genes and Genomes (KEGG) enrichment analyses further indicated that upregulated proteins in B-GEVs were enriched in pathways associated with molecular regulation, protein transport, vesicle-mediated transport, and endocytic processes (**Fig. 4f and g**), supporting the hypothesis that boiling-driven proteomic remodeling enhances vesicle internalization capacity.

**Fig. 4.**
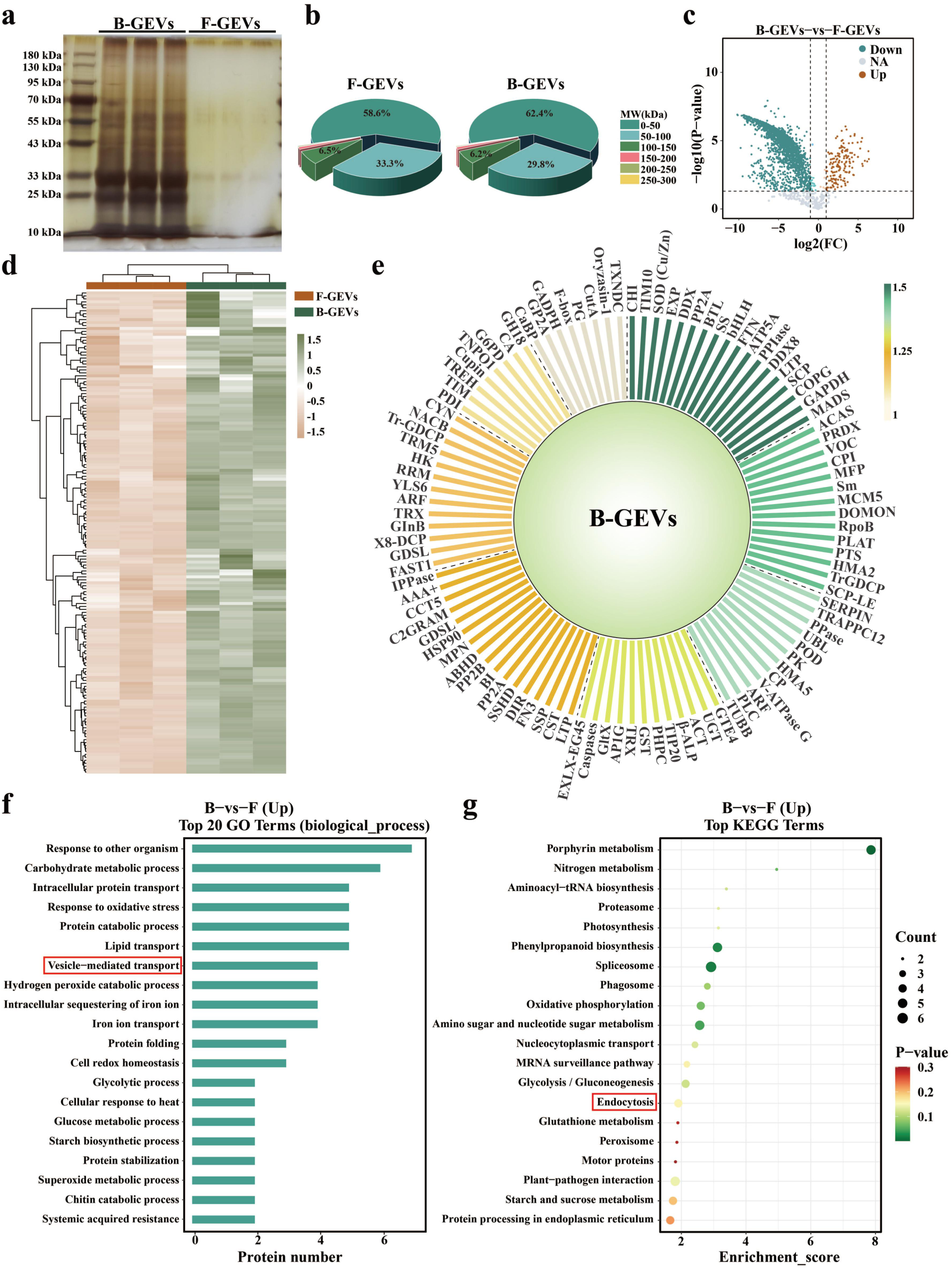
The formation of protein shell on B-GEVs was evaluated by SDS–PAGE and LC-MS/MS. **(a)**, Silver-stained gel comparing protein profiles of F-GEVs and B-GEVs. **(b)**, Protein molecular weight ratio in F-GEVs and B-GEVs. **(c)**, Volcano plot of differentially expressed proteins (|log2FC| > 1, *P* < 0.05). **(d)**, Hierarchical clustering of proteomic profiles. **(e)**, Schematic representation of upregulated B-GEVs proteins with annotated functional domains. **(f)**, GO biological process enrichment analysis of differential proteins. **(g)**, KEGG pathway mapping of protein functional networks.

### Boiling processing facilitates the incorporation of clathrin-mediated endocytosis signaling proteins into GEVs

Uptake assays both *in vivo* and *in vitro* demonstrated the crucial role of the B-GEVs protein shell in regulating uptake-related events. To further confirm the contribution of protein shell, the cellular uptake of DB-GEVs and DF-GEVs was assessed. As shown in **Fig. 5a-c**, DB-GEVs uptake levels were comparable to DF-GEVs in RAW264.7, NCM460, and HepG2 cells, supporting the role of the protein shell in mediating endocytosis. It is well established that the formation of a biomolecular corona on the surface of nanoparticles can obscure their target proteins, thereby reducing targeting efficiency (*16*). However, despite the presence of a protein shell, B-GEVs maintain their targeting function. This enhanced targeting capability is likely attributed to the unique composition of the protein shell on B-GEVs. Subcellular localization analysis revealed that the upregulated proteins in B-GEVs predominantly originated from the plasma membrane, cytoplasm, and Golgi apparatus (**Fig. 5d**), suggesting a potential role in transmembrane transport that may facilitate the efficient intracellular delivery of B-GEVs (*26*). To elucidate mechanisms underlying enhanced B-GEVs uptake, we conducted functional validation through pharmacological interrogation with pathway-specific inhibitors. InterPro analyses identified upregulation of proteins associated with clathrin-mediated endocytosis (CME) (**Fig. 5e**). Inhibitor targeting the CME pathway was utilized in the treatment of both non-phagocytic and phagocytic cells (**Fig. 5f**). The addition of chlorpromazine (CPZ), a known inhibitor of CME (*10*), remarkably decreased the cellular uptake efficiencies of B-GEVs in RAW264.7, NCM460, and HepG2 cells. Notably, a comparable internalization efficiency was observed within a 12 h period following the uptake of both B-GEVs and F-GEVs in RAW264.7 (**Fig. 5g**) and HepG2 cells (**Fig. 5h**). Moreover, the administration of CPZ resulted in a reduction in the cellular uptake efficiencies of B-GEVs compared to F-GEVs in the NCM460 cells (**Fig. 5i**). Given the limited specificity of individual inhibitors, we utilized multiple compounds targeting distinct steps of the CME pathway—namely Pitstop 2, ES9-17, and Dynole 2-24 (**Fig. S8**). The internalization of B-GEVs was consistently reduced by each inhibitor, supporting the involvement of CME in cellular uptake. Notably, despite inhibiting macropinocytosis and caveolin-mediated phagocytosis pathways, the internalization of B-GEVs was significantly enhanced (**Fig. 5g-i**). After treatment with F-GEVs or B-GEVs, the levels of membrane-associated clathrin showed no significant increase in RAW264.7, NCM460, or HepG2 cells. (**Fig. S9**). These findings further confirmed that thermal processing promotes the incorporation of CME-related proteins into B-GEVs, thereby enhancing their internalization. Overall, these findings confirm that boiling promotes the incorporation of CME-related proteins into GEVs, thereby enhancing their cellular internalization across phagocytic and non-phagocytic cells.

**Fig. 5.**
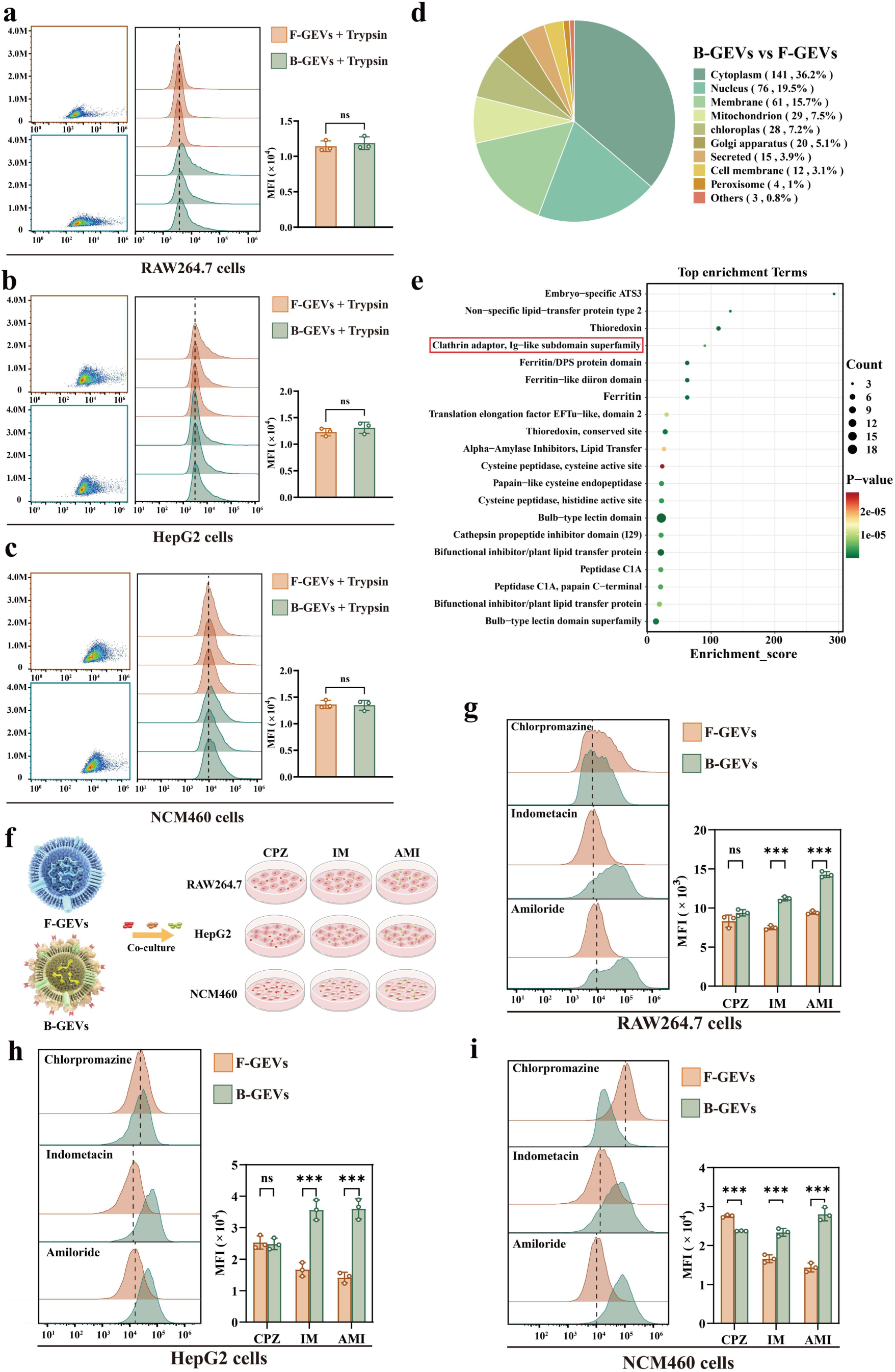
Internalization of B-GEVs is controlled by the clathrin-related endocytosis machinery. **(a-c)**, The uptake of DF-GEVs and DB-GEVs in RAW 264.7 cells (a), HepG2 cells (b), and NCM460 cells (c). n=3 biologically independent samples. The two-sided unpaired t-test method was used for statistical significance testing. ns, non-significant. **(d)**, subcellular localization analysis**. (e)**, Interpro analysis. **(f)**, Experimental design. (**g-i)**, The uptake of F-GEVs and B-GEVs in RAW 264.7 cells (g), HepG2 cells (h), and NCM460 cells (i), with the chlorpromazine (CPZ), indomethacin (IM), and amiloride (AM). n=3. The two-sided unpaired t-test method was used for statistical significance testing. *** *P* < 0.001, ns, non-significant.

### B-GEVs possess intrinsic anti-inflammatory activity as a therapeutic agent

The distinct structural and compositional features of B-GEVs compared to F-GEVs likely account for their divergent biological activities. In an LPS-stimulated RAW 264.7 macrophage model of diet-induced inflammation, B-GEVs significantly suppressed pro-inflammatory cytokines, increased anti-inflammatory IL-10, and reduced TGF-β secretion, unlike F-GEVs (**Fig. 6a and Fig. S10**). No significant differences were observed in TNF-α, IL-1β, TGF-β, and IL-6 expression between DB-GEVs and DF-GEVs treatment groups **(Fig. S11)**.

**Fig. 6.**
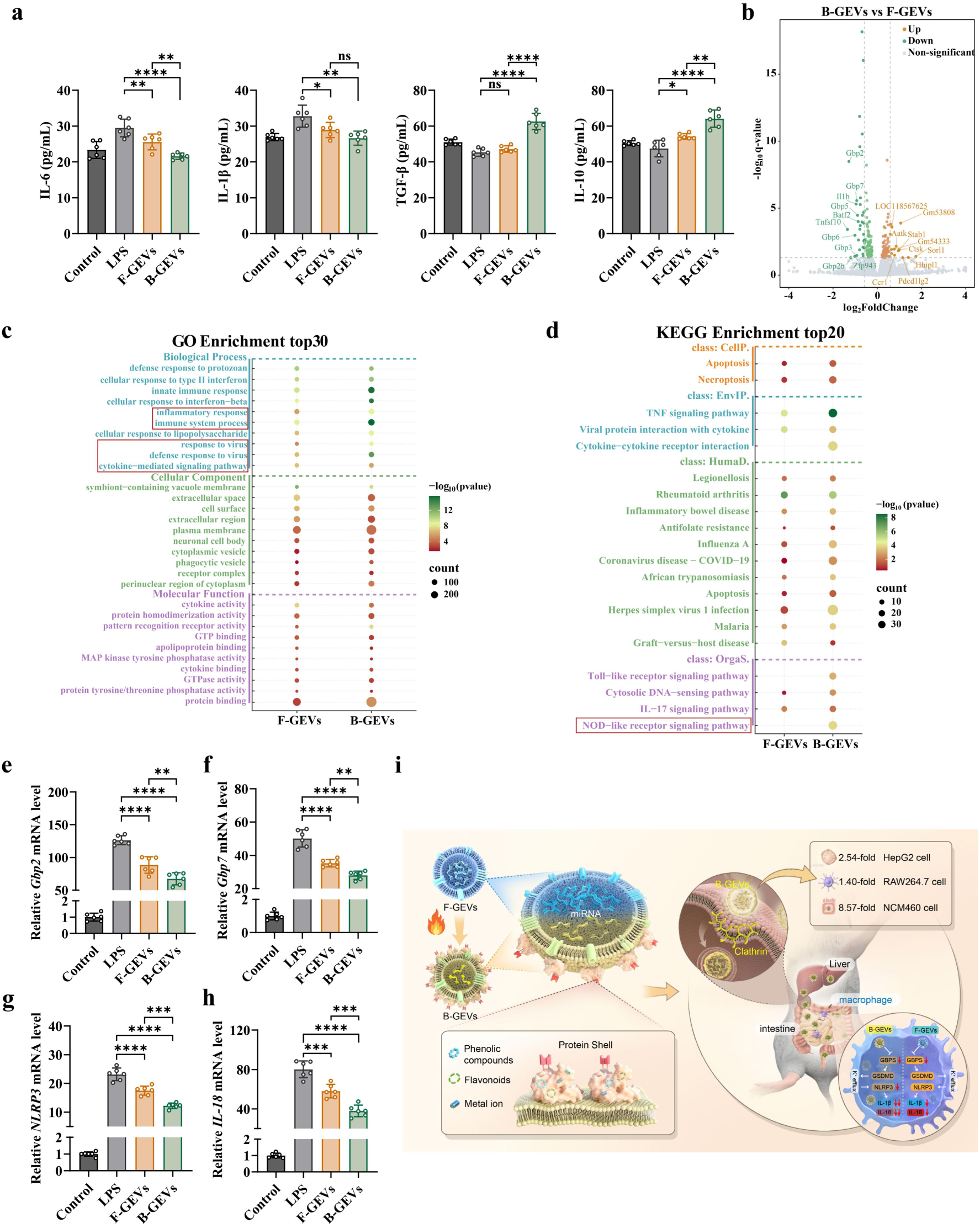
Superior anti-inflammatory performance of B-GEVs over F-GEVs. **(a)**, ELISA analysis showing the levels of inflammatory cytokines and TGF-β in macrophages transfected with F-GEVs and B-GEVs. **(b)**, Volcano plot showing differentially expressed genes between F-GEVs and B-GEVs. **(c)**, GO enrichment analysis in F-GEVs and B-GEVs-treated macrophages. **(d)**, KEGG pathway analysis. **(e-h)**, Expression of key genes in the NOD-like receptor signaling pathway after the addition of F-GEVs and B-GEVs, including *Gbp2* (e), *Gbp7* (f), *NLRP3* (g), and *IL-18* (h). **(i)**, Schematic illustrating the large-scale generation of functional B-GEVs from boiled ginger juice and their mechanism of anti-inflammatory action. All data are presented as means ± SD, n = 6. P values were calculated using two-sided one-way ANOVA post-Dunnett’s test; \**P* < 0.05, \*\**P* < 0.01, \*\*\**P* < 0.001, \*\*\*\**P* < 0.0001.

RNA-seq analyses was performed on RAW 264.7 cells treated with F-GEVs and B-GEVs to explore the transcriptomic changes induced by boiling. RNA-seq revealed distinct transcriptional profiles (**Fig. S12 and Fig. S13**), with 46 differentially expressed genes (*P* < 0.05, |log2FC| > 1) between F-GEVs and B-GEVs-treated macrophages (**Fig. S14**). Downregulation of Gbp family genes was associated with restrained inflammatory activation and a more balanced antiviral response (*27*). B-GEVs treatment significantly downregulated key inflammatory genes (i.e. *Gbp2*, *Gbp5*, *Gbp7*, *Tlr3*, and *Ifit1*) and upregulated genes associated with endocytic recycling (*Sorl1* and *Stab1*) (**Fig. 6b**). These transcriptional changes suggest that B-GEVs promote an anti-inflammatory and antiviral cellular response. The primary down-regulated biological process category involves terms like “innate immune response”, “inflammatory response”, “cellular response to interferon-beta adhesion”, “immune system process”, and “defence response to virus” (**Fig. 6c**). Notably, GO and KEGG analysis of the upregulated genes in B-GEV-treated cells indicated enrichment of pathways involved in endocytic recycling, vesicular transport, and LDL particle binding. These bioinformatic findings are consistent with our prior results (**Fig. 4f and g**) and further suggest a potential role for B-GEVs in promoting endocytic processes and facilitating active EVs uptake (**Fig. S15 and Fig. S16**). Analyses highlighted pathways linked to antiviral responses and inflammation, including NOD-like receptor, Toll-like receptor, and IL-17 signaling pathways (**Fig. 6d**). RT-qPCR validation confirmed reduced expression of *Gbp2*, *Gbp7*, *NLRP3*, and *IL-18* in B-GEVs-treated LPS-stimulated RAW264.7 macrophage (**Fig. 6e-h**), supporting the conclusion that B-GEVs exert superior anti-inflammatory effects through modulation of Gbp family genes and suppression of NLRP3 inflammasome activation. Our study demonstrates that boiling facilitates the generation of bioactive B-GEVs (**Fig. 6i**), offering a novel approach to natural engineering.

### B-GEVs as a synergistic therapeutic platform combining innate anti-inflammatory activity with efficient siRNA delivery

Given the established intrinsic anti-inflammatory properties and efficient intestinal absorption of B-GEVs, we explored their potential as an all-in-one synergistic therapy. We hypothesized that combining the innate immunomodulatory activity of B-GEVs with targeted gene silencing would improve therapeutic outcomes. To this end, we developed an oral siRNA delivery system using B-GEVs (**Fig. 7a)**. The TNF-α siRNA was loaded into both F-GEV and B-GEV using optimized electroporation parameters (300 V), achieving a loading efficiency of approximately 18% (**Fig. 7b and Fig. S17**). TEM revealed that both B-GEVs/siRNA^TNF-α^ and F-GEVs/siRNA^TNF-α^ retained their intact structures, exhibiting a typical cup-shaped morphology with nanoscale dimensions (**Fig. 7c**). Following siRNA loading, the hydrodynamic diameter increased to 146.3 ± 8.3 nm for B-GEVs/siRNA^TNF-α^ and 133.5 ± 4.5 nm for F-GEVs/siRNA^TNF-α^, while the surface potential decreased to −5.07 ± 0.77 mV and −9.70 ± 0.85 mV, respectively (**Fig. 7d and e**). Gel electrophoresis further demonstrated that the nanovesicle membranes effectively protected the encapsulated siRNA from degradation unless chemically lysed (**Fig. 7f**).

**Fig. 7.**
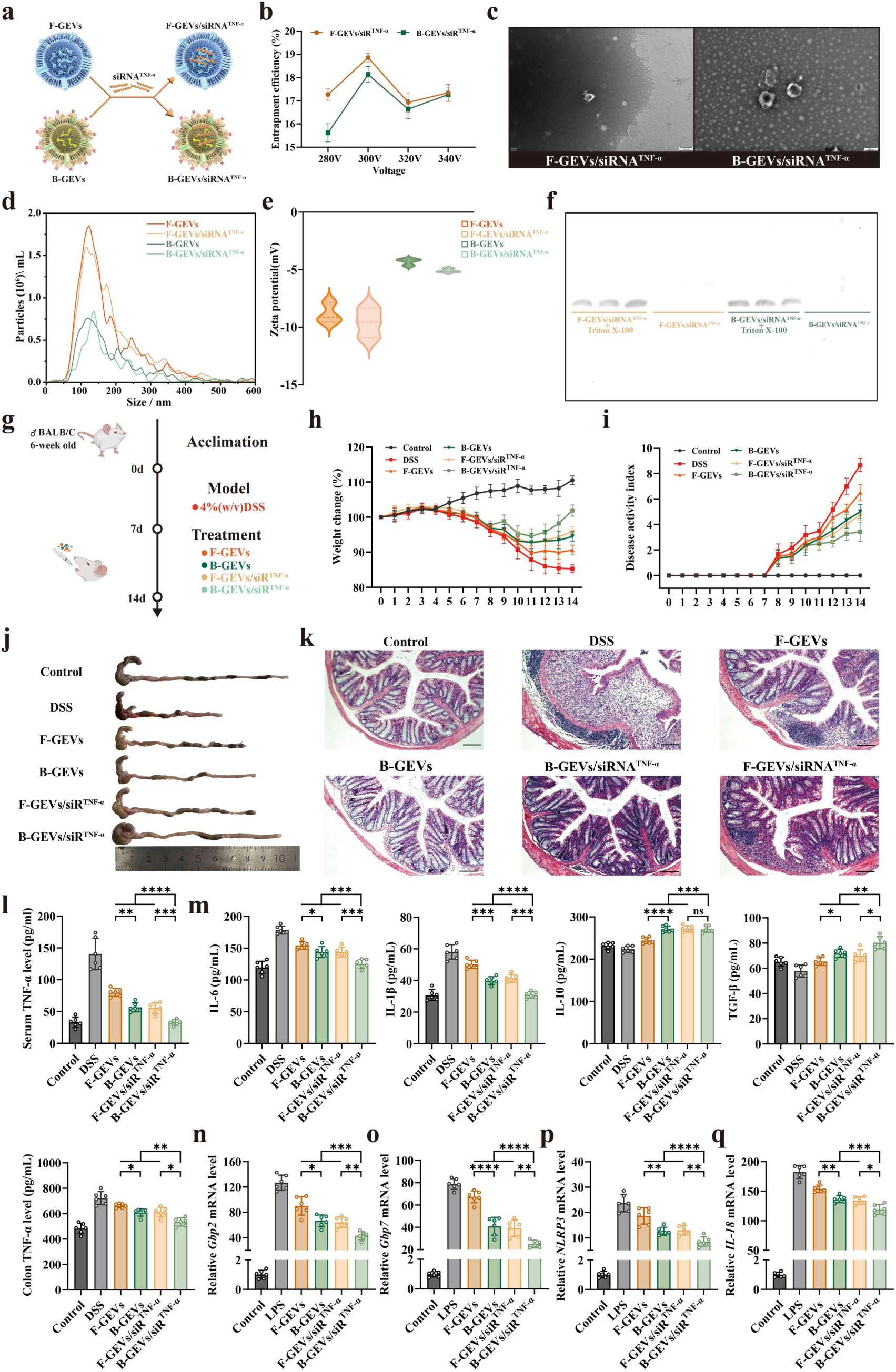
Therapeutic effects of B-GEVs/siRNA^TNF-α^ in IBD models. **(a)**, Schematic illustration of the preparation of F-GEVs/siRNA^TNF-α^ and B-GEVs/siRNA^TNF-α^. **(b)**, Loading efficiency of TNF-α siRNA into F-GEVs and B-GEVs at different electroporation voltages. **(c)**, Representative TEM photos of F-GEVs/siRNA^TNF-α^ and B-GEVs/siRNA^TNF-α^, Scale bar, 100 µm. **(d and e)**, Particle size (d) and zeta potential (e). **(f)**, RNase protection assay assessing siRNA stability. **(g)**, Experimental scheme for DSS-induced acute colitis and therapeutic administration. **(h)**, Body weight changes in each group on day 14. **(i)**, DAI score. **(j)**, Representative photographs of excised colon tissues. **(k)**, Analysis of colon tissues using H&E assays post-treatment. Scale bars: 100 μm. **(l)**, TNF-α level of serum and colon tissue after treatment of colitis. **(m)**, ELISA-based measurement of inflammatory cytokines and TGF-β levels in colon tissue. **(n-q)**, Relative mRNA expression levels of *Gbp2* (n), *Gbp7* (o), *NLRP3* (p), and *IL-18* (q) were measured by RT-qPCR. All data are presented as means ± SD, n = 6. P values were calculated using two-sided one-way ANOVA post-Dunnett’s test; \**P* < 0.05, \*\**P* < 0.01, \*\*\**P* < 0.001, \*\*\*\**P* < 0.0001, ns, non-significant.

We directly compared the therapeutic efficacy of B-GEVs/siRNA^TNF-α^, F-GEVs/siRNA^TNF-α^, F-GEVs alone, and B-GEVs alone in a DSS-induced colitis model. Mice receiving 4% DSS in drinking water began losing weight on day 5 due to severe diarrhea and reduced food intake (**Fig. 7g)**. After 6 days of treatment, the proportion of weight loss alleviated and the disease activity index (DAI) decreased following treatment with B-GEVs/siRNA^TNF-α^ or F-GEVs/siRNA^TNF-α^ compared to the DSS, F-GEVs or B-GEVs groups (**Fig. 7h and i**). Although both F-GEVs and B-GEVs treatments provided benefits, B-GEVs alone offered significantly better protection against body weight loss and colon shortening than F-GEVs (**Fig. 7h and j**), reaffirming their enhanced intrinsic activity. The synergistic effects of B-GEVs and TNF-α siRNA were evident in the significantly higher weight and lower DAI in the B-GEVs/siRNA^TNF-α^ group compared to the other groups (**Fig. 7h and i**). Treatment with B-GEVs/siRNA^TNF-α^ produced the most profound therapeutic recovery, while B-GEVs/siRNA^TNF-α^ recipients showed nearly normalizing colon length and histology compared to other groups, owing to its synergistic therapeutic effects (**Fig. 7h–j**). H&E staining indicated that the B-GEVs/siRNA^TNF-α^ group exhibited the most substantial alleviation of colonic epithelial damage (**Fig. 7k**). Furthermore, no significant histopathological alterations were observed in the liver or kidney tissues across all treatment groups (**Fig. S18**). Meanwhile, oral administration of B-GEVs/siRNA^TNF-α^ led to a significant reduction in TNF-α levels in both serum and colonic tissues, outperforming the F-GEVs, B-GEVs, and F-GEVs/siRNA^TNF-α^ groups (**Fig. 7l**). Furthermore, B-GEVs/siRNA^TNF-α^ treatment markedly decreased the pro-inflammatory cytokines IL-6 and IL-1β while elevating TGF-β (**Fig. 7m**). In contrast, it substantially reduced IL-10, a potent immunosuppressive cytokine (**Fig. 7m**). Notably, reductions in *Gbp2*, *Gbp7*, *NLRP3*, and *IL-18* were observed specifically in the B-GEVs/siRNA^TNF-α^ group compared to the other three groups (**Fig. 7n–q**). Together, these data suggest that oral administration of B-GEVs/siRNA^TNF-α^ can effectively hinder the development of intestinal inflammation, demonstrating significantly better therapeutic effects compared to B-GEVs or F-GEVs/siRNA^TNF-α^ therapy alone. The B-GEVs/siRNA^TNF-α^ platform exerts a synergistic, multi-layered anti-inflammatory effect by simultaneously modulating upstream regulatory pathways through its innate cargo and silencing downstream effector genes via delivered siRNA.

## Discussion

EVs are essential mediators of cellular communication and promising therapeutic carriers (*1*). Edible plants provide a natural source of EVs, evolved to aid nutrient absorption and inspire drug delivery strategies (*2*). Though vegetables are consumed raw or cooked, prior studies suggest cooking primarily alters phytochemical composition and physical characteristics (*3*). Ginger juice constitutes a heterogeneous system in which EVs critically contribute to its bioactivity (*4, 5*). Yet, how boiling affects the nanostructure and function of GEVs remains unclear. Our work addresses a critical oversight by demonstrating that boiling actively reconstitutes PEVs, fundamentally reframing it from a passive, potentially degradative culinary step into an active, structure-engineering strategy that enhances their health-promoting and therapeutic potential. Here, we demonstrate that thermal processing actively reconfigured B-GEVs with a protein shell enriched in native proteins, phytochemicals, and physiological metals. This reconstituted architecture not only enhances cellular uptake and anti-inflammatory efficacy but also positions B-GEVs as potent platforms for nucleic acid therapeutics.

Comparative isolation and analysis of B-GEVs and F-GEVs revealed marked differences in particle size, rigidity, and morphology. NTA indicated a moderate increase in mean diameter (∼14 nm) in polydisperse EVs samples, which is unlikely to be the primary determinant of altered biodistribution (*28*). Changes in EVs outer membrane size is associated with compositional shifts (*29*), offering a basis for differential biodistribution. Following oral administration, B-GEVs preferentially localized to intestinal and hepatic tissues—a profile pertinent to interventions targeting gut-liver axis disorders. Because efficient cellular internalization is essential for PEVs to exert cross-kingdom effects (*10*, *30*, *31*), we investigated the basis for the enhanced uptake of B-GEVs. The formation of protein shells in B-GEVs was confirmed via comparative analysis of morphology, protein concentration, particle size, and rigidity relative to F-GEVs. As in biological systems that undergo stimulus-induced structural reconfiguration in liquid environments (*18*, *19*), the protein shell in B-GEVs appears to arise from thermally driven interfacial self-assembly of ginger-derived proteins, phytochemicals, and metal ions. Meanwhile, miRNA loss was observed in B-GEVs following boiling, suggesting selective cargo stabilization. The results suggest that the nano-structural reconstitution of B-GEVs through boiling represents a scalable strategy for function-oriented food engineering.

Our proteomic analyses revealed substantial remodeling of the B-GEVs proteome, providing mechanistic insights into their enhanced uptake properties. Proteomic profiling revealed that the B-GEVs shell is enriched in proteins associated with vesicular trafficking and endocytosis, including ARF1 and adaptor complex components essential for clathrin-coated vesicle formation (*32*, *33*). Given that endocytosis is essential for intracellular delivery of EVs cargo (*34*, *35*), our data indicating elevated internalization of B-GEVs by phagocytes, hepatocytes, and enterocytes supports the notion that boiling-induced proteomic changes facilitate uptake. Most notably, this reconstituted surface architecture mediates a striking 8.57-fold increase in uptake efficiency specifically within intestinal epithelial cells. EVs uptake generally proceeds via macropinocytosis, phagocytosis, and clathrin-/caveolin-mediated endocytosis (*36*). Engineered EVs carrying fusion proteins can facilitate targeted uptake by recipient cells (*37*, *38*). Our proteomic data indicated an upregulation of proteins related to transport and clathrin-mediated endocytosis, suggesting that enhanced uptake of B-GEVs is CME-dependent. Uptake assays both *in vivo* and *in vitro* demonstrated the crucial role of the B-GEVs protein shell in regulating uptake-related events. The observed enrichment of CME-related proteins in B-GEVs supports these findings. The surface of B-GEVs thus appears to actively engage host cell’s endocytic machinery—a feature typically engineered into synthetic nanoparticles but here emerging from simple thermal processing.

Ginger and its extracts have a long history of medicinal use, particularly in alleviating gastrointestinal disorders (*5*, *8*, *9*, *10*, *11*). We investigated whether boiling—a common culinary practice—could enhance the bioactivity of GEVs. Previous studies have established that edible GEVs exhibit immunomodulatory properties (*9*, *39*). Our findings demonstrate that thermal processing substantially enhances the inflammatory functionality of GEVs, suggesting practical applications for B-GEVs as bioactive components in dietary interventions. Specifically, we found that B-GEVs exert a pronounced inhibitory effect on the NLRP3 inflammasome which is a central mediator of inflammatory pathology. Mechanistically, this suppression is linked to their ability to modulate the expression of Gbp family genes, a class of interferon-inducible GTPases that critically regulate NOD-like receptor signaling and inflammasome activation (*40*). By attenuating this key inflammatory cascade, B-GEVs effectively dampen NOD-like receptor-mediated pro-inflammatory signaling. In a DSS-induced colitis model, B-GEVs effectively attenuated inflammatory signaling, demonstrating that a simple thermal step can transform dietary EVs into potent, physiologically relevant immunomodulators.

We developed B-GEVs into an oral siRNA delivery platform by exploiting their enhanced intestinal uptake and intrinsic tissue tropism. Our work redefines B-GEVs from versatile nanoparticles to a programmable, synergistic therapeutic platform. In IBD, where single-target therapies often show limited efficacy (*41*), B-GEVs/siRNA^TNF-α^ embodies an integrated synergistic strategy. The native components of B-GEVs temper upstream inflammatory pathways, while the siRNA payload directly silences a downstream mediator like TNF-α. It also suggests that the platform’s utility can be extended by loading different siRNA cargos to target other disease-specific genes, all while leveraging the consistent foundational immunomodulation and targeting provided by the B-GEVs scaffold.

## Materials and Methods

### Materials

The BCA protein assay kit, LPS, 3,3′-dioctadecyloxacarbocyanine perchlorate (DIO), and 6-indolecarbamidine dihydrochloride (DAPI) were procured from Shanghai Beyotime Biotechnology. The exosome protein specific lysis buffer and 1,1′-dioctadecyl-3,3,3′,3′-tetramethylindotricarbocyanine iodide (DIR) were purchased from Shanghai Nonin Biological Technology Co., Ltd. ELISA kits were supplied by Shanghai Youxuan Biotechnology Co., Ltd. Fetal Bovine Serum (FBS), amiloride, indomethacin, and chlorpromazine were supplied by Merck & Co., Inc., while Dulbecco’s modified Eagle’s medium (DMEM) was provided by Thermo Fisher Scientific Inc. The CCK-8 assay kit was purchased from Hefei Biosharp. Penicillin and streptomycin were obtained from Invitrogen. All other chemicals used were of analytical grade.

### Isolation and purification of F-GEVs and B-GEVs

The fresh ginger rhizomes were thoroughly washed with ultrapure water, then sliced into small pieces using a knife and placed into a blender. After being squeezed, the juice was immediately filtered through a 100-mesh sieve to remove residual fibers. Ginger juice from the same batch was divided into two groups: untreated sets and boiled-treated sets. The boiled-treated group was boiled at 100 °C for 60 minutes, while the untreated group was stored at 4 °C. All samples were stored at 4 °C prior to EVs isolation. F-GEVs and B-GEVs were isolated using differential centrifugation and commercial exosome isolation kits as previously described (*13*, *20*). Purified EVs were kept at -80 °C for future use. Protein concentration of F-GEVs and B-GEVs was measured using a BCA protein assay kit.

### Characterization of F-GEVs and B-GEVs

The morphology of F-GEVs and B-GEVs was observed using a HT7700 transmission electron microscope (TEM). Briefly, samples were placed onto 200-mesh copper grids for 5 min, negatively stained with phosphotungstic acid for 1 minute, and imaged at an accelerating voltage of 100 kV. Particle counts and sizes were determined by NTA (M Particle Metrix, Meerbusch, Germany), and the zeta potential was measured using laser diffraction spectrometry (Malvern Mastersizer 2000, Malvern).

### Cytotoxicity assay

RAW 264.7 and NCM460 cells were plated in 96-well plates at a density of 5 × 10^3^ cells per well and cultured for 24 h. Cells were then treated with serial dilutions of F-GEVs or B-GEVs (ranging from 0 to 5×10^10^ particles/mL) for 24 h. Cell viability was assessed using the CCK-8 assay, and absorbance was measured at 450 nm.

### Determination of total phenols and total flavonoids

A sample solution was prepared for total phenols and total flavonoids content analysis by homogenizing the GEVs extract with 50% ethanol. For total phenolic content analysis, 40 μL of the sample was mixed with 1 mL of Folin-Ciocalteu reagent. After 5 min, 0.8 mL of 7.5% sodium carbonate was added. The mixture was incubated in the dark for 30 minutes, and absorbance was measured at 765 nm. For total flavonoids, 1 mL of the sample was mixed with 150 μL 5% sodium nitrite, followed by 300 μL 10% aluminum nitrate after 5 minutes, then 1 mL of 1 M sodium hydroxide. The final volume was adjusted to 4 mL with deionized water. Absorbance was recorded at 510 nm.

### Metal analysis

The concentrations of copper, iron, zinc, manganese, magnesium, potassium, and calcium were quantified using inductively coupled plasma mass spectrometry (ICP-MS). Lyophilized GEVs samples were digested overnight at room temperature in 50 μL of 65% (v/v) nitric acid. After heating to 90 °C for 20 minutes, samples were diluted to 0.5 mL with 1% (v/v) nitric acid and analyzed using an Agilent 7900 ICP-MS with helium collision mode under standard multi-element settings.

### Dynamic simulation of protein layer formation

Ginger-derived proteins were co-incubated with F-GEVs at 100 °C. At specified time points, protein concentrations were measured using a BCA assay. Proteins were pre-labeled with Alexa Fluor 488 and incubated with F-GEVs at 100 °C for 60 min. Following incubation, F-GEVs were counterstained with DiD. After ultrafiltration to remove excess dye, DiD-labeled F-GEVs were imaged using confocal laser scanning microscopy (Olympus FV4000). Fluorescence intensity was subsequently measured using a microplate reader (SuPerMax 3200) at excitation wavelengths of 488 nm.

### Proteomics analysis of F-GEVs and B-GEVs

Protein extraction from F-GEVs and B-GEVs was conducted using Exosome Protein Specific Lysis Buffer. The subsequent proteomic data analysis was executed by Shanghai OE Biotech Co., Ltd. (Shanghai, China). LC-MS/MS was performed on a Tims TOF Pro mass spectrometer (Bruker) with an Easyspray ion source. Proteins were digested with trypsin, desalted, and analyzed using data-independent acquisition (DIA). MS/MS spectra were analyzed using Spectronaut Pulsar 18.7 (Biognosys), with searches performed against the Uniprot database (uniprot_proteome_UP000734854_Zingiber officinale_20240821.fasta).

### Rigidity measurements of F-GEVs and B-GEVs

10 μL of F-GEVs and B-GEVs were adsorbed onto clean mica sheets, then subjected to multiple rinses with an appropriate volume of pure water. Following the removal of excess liquid using filter paper, the mica sheets were fixed onto an iron plate. Subsequently, the differences in surface rigidity among the F-GEVs and B-GEVs were analyzed using Dimension Icon.

### Secondary structure analysis of ginger proteins pre- and post-boiling

Ginger proteins were dissolved in phosphate buffer (10 mM, pH 7.4) to a final concentration of 0.2 mg·mL⁻¹. Fresh samples were analyzed directly, while boiled samples were obtained by heating the solution at 100 °C for 60 min and then rapidly cooling it to room temperature. To remove insoluble aggregates, the samples were centrifuged at 12,000 × g for 10 min, and the supernatants were collected for analysis. Far-UV CD spectra were recorded at 25 °C using a spectropolarimeter equipped with a 0.1 cm path length quartz cuvette. Data were collected from 190 to 320 nm at 100 nm/min with a 1 nm bandwidth and 1 s response time. Each reported spectrum is the average of three consecutive scans, corrected against a buffer baseline, and expressed as mean residue ellipticity (mdeg).

### ITC analysis

We used isothermal titration calorimetry (MicroCal PEAQ-ITC) to determine the binding affinity of F-GEVs for ginger proteins before and after boiling. GEVs (1.3 × 10^10^ particles/ mL) were loaded into the sample cell, and ginger protein (2 mg/mL) was placed in the injection syringe. Titrations were carried out at 25 °C with stirring at 750 rpm. The protocol included 19 injections: an initial 0.4 μL injection followed by 18 injections of 2 μL each, with 120 s intervals. Data analysis was performed with the instrument’s dedicated software.

### SDS-PAGE

Equal numbers of particles from F-GEVs and B-GEVs were lysed and loaded onto 12% SDS–PAGE gels. Protein ladders (Precision Plus Dual Color, BioRad) were used for molecular weight estimation. Gels were electrophoresed at 80 V for 30 min, then at 120 V for 60 min. Bands were visualized via silver staining and imaged using a Tanon MINI Space 3000 system.

### Cellular uptake assay

RAW 264.7 cells were cultured on confocal dishes and incubated with DiR-labeled F-GEVs or B-GEVs (5×10^9^ particles/mL) for 4 or 12 h. After incubation, cells were washed with cold PBS, fixed with 4% paraformaldehyde, and stained with DAPI and DiO. Uptake was visualized using a Leica STELLARIS5 confocal microscope. No significant difference in DiO fluorescence intensity was observed between F-GEVs and B-GEVs (**Fig. S19**).

To further confirm the uptake of F-GEVs or B-GEVs *in vitro*, a quantitative analysis via flow cytometry was conducted. The F-GEVs or B-GEVs were labeled with DiO. Utilizing a flow cytometer, fluorescence intensities (λ = 484 nm) were measured in RAW264.7 cells, HepG2 cells, and NCM460 cells, thereby enabling a clearer comparison of the uptake variations across the various experimental groups.

### Uptake inhibition studies

RAW 264.7, HepG2, and NCM460 cells were seeded in 12-well plates (1×10⁵ cells/well) and cultured for 24 h. Cells were pretreated for 1 h with the following inhibitors: amiloride, indomethacin, Pitstop 2, ES9-17, Dynole 2-24, or chlorpromazine. Subsequently, F-GEVs or B-GEVs were added and incubated with the cells for 12 h. Cellular uptake of GEVs was then assessed by flow cytometry to quantify internalization efficiency under each condition. To investigate the contribution of surface proteins, F-GEVs and B-GEVs were subjected to digestion with 0.25% trypsin at a protein-to-enzyme ratio of 2:1 for 1 h at 37 °C. After removing trypsin via ultrafiltration, the resulting particles were designated as DF-GEVs and DB-GEVs, respectively. The fold change in uptake was calculated directly based on the mean fluorescence intensity (MFI) of the reference control using the following formula:

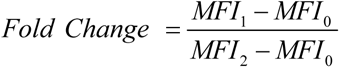

where MFI_1_, MFI_2_, and MFI_0_ represent the MFI values of the B-GEVs, F-GEVs, and control group, respectively.

### In *vivo* distribution of F-GEVs and B-GEVs

Six-week-old male C57BL/6 mice (body weight 20 ± 2 g) were obtained from Hefei Qingyuan Biotechnology Co., Ltd. After a week of cohabitation, the mice were divided into two groups, with three replicates per time point. Each group received an oral administration of EVs at a concentration of 5×10^9^ particles. The distribution of EVs within the body was subsequently monitored using an in vivo imaging system (IVIS; PerkinElmer, Waltham, USA) at 1, 4, 6, 10, 18, and 24 h post-administration. At each time point, mice from the corresponding group were euthanized, and major organs were excised for ex vivo imaging with IVIS. All animal experiments were approved by the Institutional Animal Care and Use Committee of Hefei University of Technology (Protocol No. HFUT20240924001).

### MiRNA detection

The expression levels of several specific miRNAs (cca-miR156b, gma-miR-6300, zma-miR164h-5p, osa-miR-164c, gma-miR396a-3p, aly-miR396a-5p, osa-miR164d, aly-miR159a-3p, and vvi-miR396a) quantified by RT-qPCR. miRNA sequence information was shown in **Table S2**. The samples were reverse transcribed using the miRNA first strand cDNA synthesis kit, and 10 pM of synthetic cel-miR-39-3p was added to each sample as a quantitative. Relative expression was calculated using the 2^-ΔΔCt^ method. Primer sequences were shown in **Table S3**, and the downstream primer sequence was CGCCCGCCCGCTCCCAAGAT.

### RNA-seq

RNA-seq was performed by OE Biotech Co., Ltd. Libraries were sequenced on an Illumina NovaSeq 6000 platform (150 bp paired-end). Raw reads were filtered with fastp, aligned to the reference genome using HISAT2, and expression levels quantified with HTSeq-count. Differential expression analysis was conducted with DESeq2 (Q < 0.05 and fold change > 2 or < 0.5 as thresholds for significance). GO and KEGG enrichment analyses were performed to characterize DEGs.

### Preparation of F-GEVs/siRNA^TNF-α^ and F-GEVs/siRNA^TNF-α^

Exosomes were loaded with siRNA via electroporation using the Gene Pulser Xcell system (Bio-Rad). Briefly, exosome-siRNA mixtures were transferred into 0.4 cm gap cuvettes and electroporated at varying voltages (280 V, 300 V, 320 V, and 340 V) with a capacitance of 125 μF, resulting in pulse durations of 5–10 ms. Following the pulse, samples were incubated on ice for a minimum of 120 min to facilitate membrane recovery. Unincorporated siRNA was subsequently removed by ultrafiltration using 100 kDa Amicon Ultra-0.5 centrifugal filter units at 14,000 × g for 15 min.

### Stability assessment of F-GEVs/siRNA^TNF-α^ and B-GEVs/siRNA^TNF-α^

To evaluate the encapsulation efficiency and integrity of siRNA loaded into F-GEVs and B-GEVs, denaturing polyacrylamide gel electrophoresis (PAGE) was performed. Samples were prepared under two conditions: intact GEVs/siRNA^TNF-α^ and those treated with Triton X-100 to disrupt vesicular membranes. The samples were resolved on a 12% urea-polyacrylamide gel at 110 V for 30 min. Post-electrophoresis, the gel was imaged using a Tanon MINI Space 3000 system.

### Evaluation of therapeutic effects

BALB/c mice were randomly allocated into six groups: (1) Healthy control (PBS); (2) DSS model (4.0% DSS); (3) DSS + F-GEVs (5×10^9^ particles); (4) DSS + B-GEVs (5×10^9^particles); (5) DSS + F-GEVs/siRNA^TNF-α^ (30 nmol/kg siRNA); and (6) DSS + B-GEVs/siRNA^TNF-α^ (30 nmol/kg siRNA). To evaluate the severity of colitis, the DAI was calculated as the summation of scores for body weight loss, stool consistency, and rectal bleeding (scale 0-4). Mice were euthanized on day 14, at which point colons and major organs were harvested for histopathological analysis. Colon, kidney, and liver tissues were fixed in formalin, embedded in paraffin, and subsequently stained with hematoxylin and eosin (H&E). All animal experiments were approved by the Institutional Animal Care and Use Committee of Hefei University of Technology (Protocol No. HFUT20240924001).

### Quantitative real-time polymerase chain reaction (RT-qPCR)

Total RNA was extracted using TRIzol reagent following the manufacturer’s protocol, reverse transcribed, and quantified using standard RT-qPCR procedures. Primer sequences are provided in **Table S4**.

### Statistics

The experimental results were performed using GraphPad Prism, with values expressed as mean ± standard deviation (SD). Data were subjected to one-way ANOVA analysis, and statistical significance was determined at * *P* < 0.05, ** *P* < 0.01, *** *P* < 0.001, **** *P* < 0.0001, with ns indicating no significance.

## Acknowledgments

This work was supported by the National Natural Science Foundation of China (32102128), the Natural Science Foundation of Anhui Province (2508085MC046), and the Fundamental Research Funds for the Central Universities (PA2024GDSK0094), and the Key R&D Program of Anhui Province (2023n06020010)

## Author contributions

L.Z. and L.Y. conceived and supervised the study. L.H. planned and performed the research. L.H. and J.C. contributed to the statistical analysis design and execution. S.G., M.L., Z.Z., and X.W. analyzed the data. H.H., C.L., and Y.M. contributed to the experimental design and provided critical feedback on the manuscript. L.H. and L.Y. wrote the manuscript. All authors discussed the results, analyzed the data, and contributed to the preparation and editing of the manuscript.

## Data and materials availability

All data needed to evaluate the conclusions in the paper are present in the paper and/or the Supplementary Materials. Additional data related to this paper may be requested from the authors.

## References

1. Ripoll, L. et al. Biology and therapeutic potential of extracellular vesicle targeting and uptake. Nat. Rev. Mol. Cell Biol. (2026).

2. Deng, Z. et al. Broccoli-derived nanoparticle inhibits mouse colitis by activating dendritic cell AMP-activated protein kinase. Mol. Ther. 25(7), 1641–1654 (2017).

3. Moltedo, A. et al. Raw versus cooked food matching: Nutrient intake using the 2015/16 Kenya Integrated Household Budget Survey. J. Food Compos Anal. 102, 103879 (2021).

4. Xu, L, et al. A review of fruit juice authenticity assessments: Targeted and untargeted analyses. Crit Rev Food Sci Nutr. 62(22):6081–6102(2022).

5. Zhuang, X. et al. Ginger-derived nanoparticles protect against alcohol-induced liver damage. J. Extracell. Vesicles. 4, 28713 (2015).

6. Narra, F. et al. Impact of thermal processing on polyphenols, carotenoids, glucosinolates, and ascorbic acid in fruit and vegetables and their cardiovascular benefits. Compr. Rev. Food Sci. Food Saf. 23(6), e13426 (2024).

7. Carmody, R. N. et al. Cooking shapes the structure and function of the gut microbiome. Nat. Microbiol. 4(12), 2052–2063 (2019).

8. Naz, S. et al. Medicinal uses and bioactivities of ginger – a detailed review. Int. J. Chem. Biochem. Sci. 8, 71–77 (2015).

9. Yan, L. et al. Ginger exosome-like nanoparticle-derived miRNA therapeutics: A strategic inhibitor of intestinal inflammation. J. Adv. Res. 69, 1–15 (2024).

10. Teng, Y. et al. Plant-derived exosomal microRNAs shape the gut microbiota. Cell Host Microbe. 24, 637–652 (2018).

11. Wang, X. et al. An abundant ginger compound furanodienone alleviates gut inflammation via the xenobiotic nuclear receptor PXR in mice. Nat Commun. 16(1), 1280 (2025).

12. Nishidono, Y. & Tanaka, K. Effect of drying and processing on diterpenes and other chemical constituents of ginger. J Nat Med. 77, 118–127 (2023).

13. Wang, Q. et al. Delivery of therapeutic agents by nanoparticles made of grapefruit-derived lipids. Nat. Commun. 4, 1867 (2013).

14. Sundaram, K. et al. Garlic exosome-like nanoparticles reverse high-fat diet induced obesity via the gut/brain axis. Theranostics. 12, 1220–1246 (2022).

15. Li, C. et al. Surface-associated proteins on extracellular vesicles remodel the tumor microenvironment by potentiating TGF-β signaling in a contact-dependent manner. Adv. Sci. e13286 (2025).

16. Liam-Or R. et al. Cellular uptake and in vivo distribution of mesenchymal-stem-cell-derived extracellular vesicles are protein corona dependent. Nat. Nanotechnol. 19, 846–855 (2024).

17. Li, C. et al. Therapeutic hypothermia reprograms nanocarrier protein corona via apolipoprotein C1 enrichment for precision cardiovascular therapy. Proc. Natl. Acad. Sci. U.S.A. 122, e2515072122 (2025).

18. Sahtoe, D. D. et al. Reconfigurable asymmetric protein assemblies through implicit negative design. Science. 375, eabj7662 (2022).

19. Quiles, J. M. & Gustafsson, Å. B. The role of mitochondrial fission in cardiovascular health and disease. Nat. Rev. Cardiol. 19, 723–736 (2022)

20. Xu, X. H. et al. Plant exosomes as novel nanoplatforms for microRNA transfer stimulate neural differentiation of stem cells in vitro and in vivo. Nano Lett. 21, 8151–8159 (2021).

21. Yin, L. F. et al. Characterization of the microRNA profile of ginger exosome-like nanoparticles and their anti-inflammatory effects in intestinal Caco-2 cells. J. Agric. Food Chem. 70, 4725–4734 (2022).

22. Xu, F. et al. Restoring oat nanoparticles mediated brain memory function of mice fed alcohol by sorting inflammatory Dectin-1 complex into microglial exosomes. Small. 18, e2105385 (2022).

23. Dad, H. A. et al. Plant exosome-like nanovesicles: Emerging therapeutics and drug delivery nanoplatforms. Mol. Ther. 29, 13–31 (2021).

24. Xiang, J. et al. Super-self-assembly extraction from natural herbs. Nano Res. 18, 949–958 (2025).

25. Xu, F. et al. Using wool keratin derived metallo-nanozymes as a robust antioxidant catalyst to scavenge reactive oxygen species generated by smoking. Small. 18, e2201205 (2022).

26. Garcia-Martin, R. et al. Tissue differences in the exosomal/small extracellular vesicle proteome and their potential as indicators of altered tissue metabolism. Cell Rep. 38, 110277 (2022).

27. Zhang, Rongzhao et al. When human guanylate-binding proteins meet viral infections. J Biomed Sci. 28(1), 17 (2021).

28. Walkey C. D. et al. Nanoparticle size and surface chemistry determine serum protein adsorption and macrophage uptake. J. Am. Chem. Soc. 134, 2139–2147 (2012).

29. Turner, L. et al. Helicobacter pylori outer membrane vesicle size determines their mechanisms of host cell entry and protein content. Front. Immunol. 9, 1466 (2018).

30. Teng, Y. et al. Plant-derived exosomal microRNAs inhibit lung inflammation induced by exosomes SARS-CoV-2 Nsp12. Mol. Ther. 29, 2424–2440 (2021).

31. Zhang, L. et al. Lemon-derived extracellular vesicle-like nanoparticles block the progression of kidney stones by antagonizing endoplasmic reticulum stress in renal tubular cells. Nano Lett. 23, 1555–1563 (2023).

32. Doray, B. & Kornfeld, S. Gamma subunit of the AP-1 adaptor complex binds clathrin: Implications for cooperative binding in coated vesicle assembly. Mol. Biol. Cell. 12, 1925–1935 (2001).

33. Mathieu, M. et al. Specificities of secretion and uptake of exosomes and other extracellular vesicles for cell-to-cell communication. Nat. Cell Biol. 21, 9–17 (2019).

34. Zhao, B. et al. Exosome-like nanoparticles derived from fruits, vegetables, and herbs: Innovative strategies of therapeutic and drug delivery. Theranostics. 14, 4598–4621 (2024).

35. D’Souza-Schorey, C. & Chavrier, P. ARF proteins: Roles in membrane traffic and beyond. Nat. Rev. Mol. Cell Biol. 7, 347–358 (2006).

36. Li, Z. et al. Fusion protein engineered exosomes for targeted degradation of specific RNAs in lysosomes: A proof-of-concept study. J. Extracell. Vesicles. 9, 1816710 (2020).

37. Xie, Q. et al. Cellular uptake of engineered extracellular vesicles: Biomechanisms, engineered strategies, and disease treatment. Adv. Healthc. Mater. 13, e2302280 (2024).

38. Kumar, A. et al. Ginger nanoparticles mediated induction of Foxa2 prevents high-fat diet-induced insulin resistance. Theranostics. 12, 1388–1403 (2022).

39. Chen, X. et al. Similarly, exosome-like nanoparticles from ginger rhizomes inhibited NLRP3 inflammasome activation. Mol Pharm. 16(6):2690–2699 (2019).

40. Meunier, E., Wallet, P., Dreier, R. et al. Guanylate-binding proteins promote activation of the AIM2 inflammasome during infection with Francisella novicida. Nat Immunol. 16, 476–484 (2015).

41. Neurath MF. Targeting immune cell circuits and trafficking in inflammatory bowel disease. Nat Immunol. 20(8):970–979 (2019).

