## Supplementary for "Thermal Nano-Engineering of Ginger Extracellular Vesicles for Targeted Oral Therapy of Colitis"

**Supplementary Figure legends**

**Supplementary Figures**

Supplementary Figure 1. Cell viability of RAW264.7, NCM460, and HepG2 cells after 24 h treatment with F-GEVs and in B-GEVs.

Supplementary Figure 2. Flow cytometry gating strategy for analysis of F-GEVs and B-GEVs uptake in HepG2, NCM460, and RAW264.7 cells.

Supplementary Figure 3. Uptake of B-GEVs or F-GEVs in RAW264.7 cells detected by confocal microscopy (a-b). Blue: nucleus; Green: cell membrane; Red: F-GEVs or B-GEVs.

Supplementary Figure 4. The particle counts of F-GEVs and B-GEVs determined by NTA.

Supplementary Figure 5. Relative levels of selected miRNAs found in B-GEVs after boiling.

Supplementary Figure 6. Mass error analysis and correlation analysis of the DAI method based on the LC-MS/MS proteomics workflow.

Supplementary Figure 7. Venn diagram comparing significantly altered proteins as identified.

Supplementary Figure 8. Cellular uptake of B-GEVs under different inhibitors of clathrin-mediated endocytosis.

Supplementary Figure 9. Immunofluorescence analysis of membrane-associated clathrin expression in HepG2 cells, NCM460 cells, and RAW264.7 cells.

Supplementary Figure 10. The mRNA expression levels of *IL-6*, *IL-1β*, *TGF-β*, and *IL-10* were measured by RT-qPCR.

Supplementary Figure 11. Anti-inflammatory effect of DF-GEVs and DB-GEVs *in vitro*.

Supplementary Figure 12. Evaluation of biological replicates and inter-sample correlations.

Supplementary Figure 13. Venn diagrams visually illustrated the shared and unique differentially expressed genes across the comparison groups.

Supplementary Figure 14. Heat map of key gene expression showing differentially expressed genes between F-GEVs and B-GEVs.

Supplementary Figure 15. Downregulation of inflammatory genes in the B-GEVs treatment group.

Supplementary Figure 16. GO and KEGG pathway enrichment analyses of up-regulated genes in B-GEVs.

Supplementary Figure 17. RNA gel assay for determining siRNA encapsulation efficiency in F-GEVs/siRNA^TNF-α^ and B-GEVs/siRNA^TNF-α^.

Supplementary Figure 18. Representative H&E-stained histological sections of kidney and liver tissues.

Supplementary Figure 19. Quantitative analysis of fluorescence intensity from DIO-labeled F-GEVs and B-GEVs.

**Supplementary Tables**

Supplementary Table 1. Proteomic profile of significantly up-regulated proteins in B-GEVs compared to F-GEVs.

Supplementary Table 2. Sequence of miRNA mimics.

Supplementary Table 3. Primer sequence of the GEVs-derived miRNA.

Supplementary Table 4. Primer sequences for RT-qPCR.

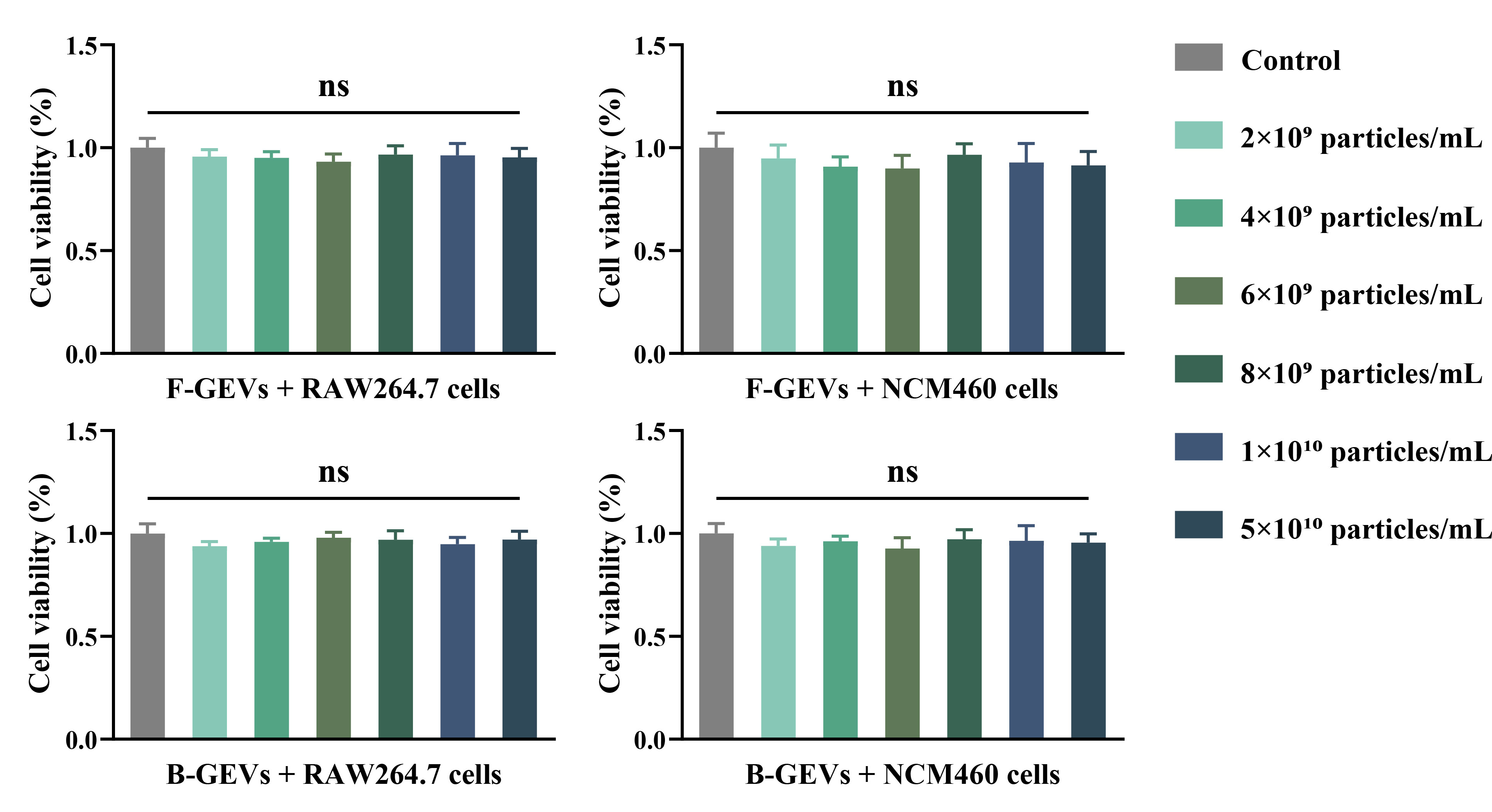

**Figure S1. Cell viability of RAW264.7, NCM460, and HepG2 cells after 24 h treatment with F-GEVs and B-GEVs.** All data are presented as means ± SD, n = 6. P values were calculated using two-sided one-way ANOVA post-Dunnett’s test; ns, non-significant.

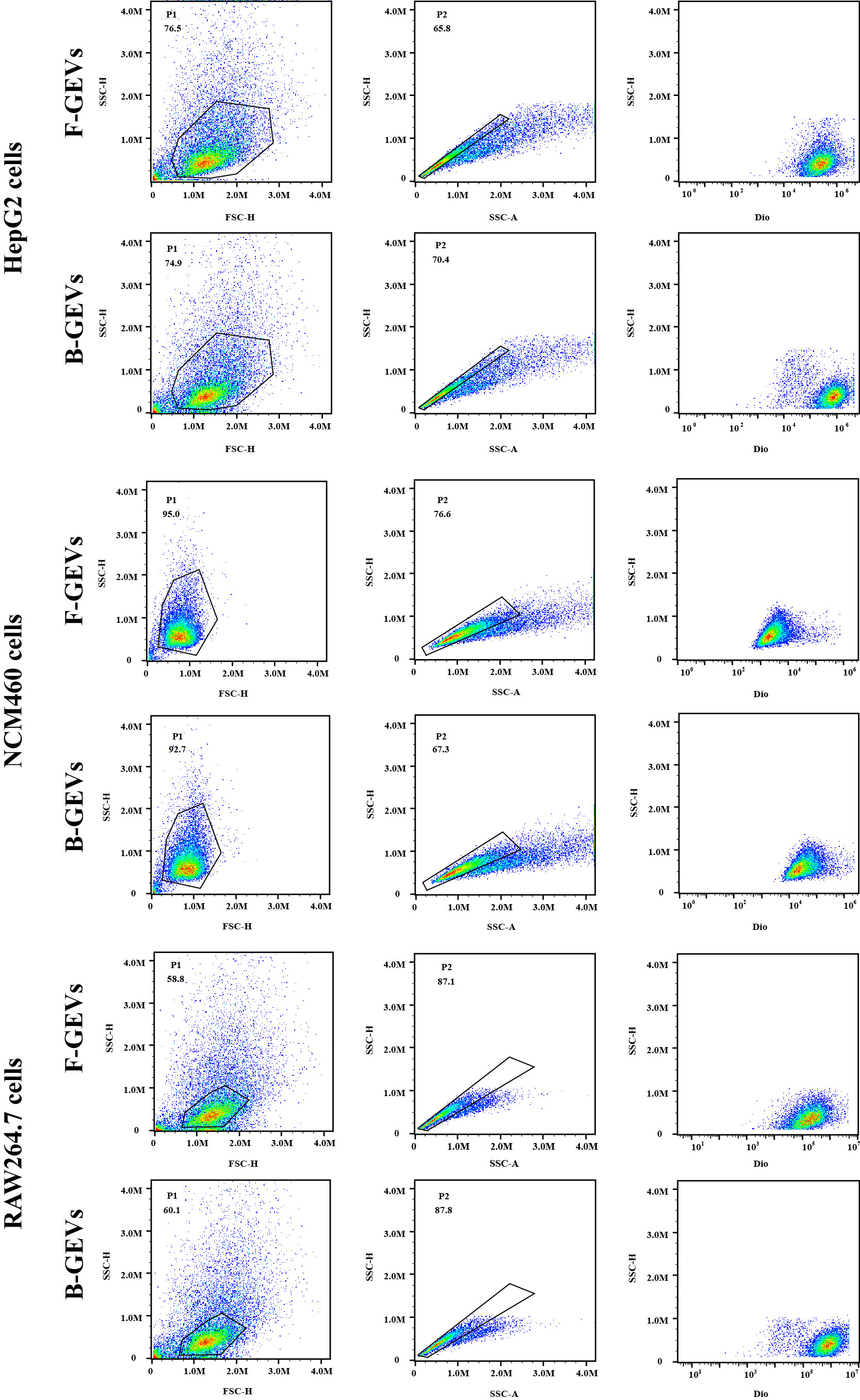

**Figure S2. Flow cytometry gating strategy for analysis of F-GEVs and B-GEVs uptake in HepG2, NCM460, and RAW264.7 cells.**

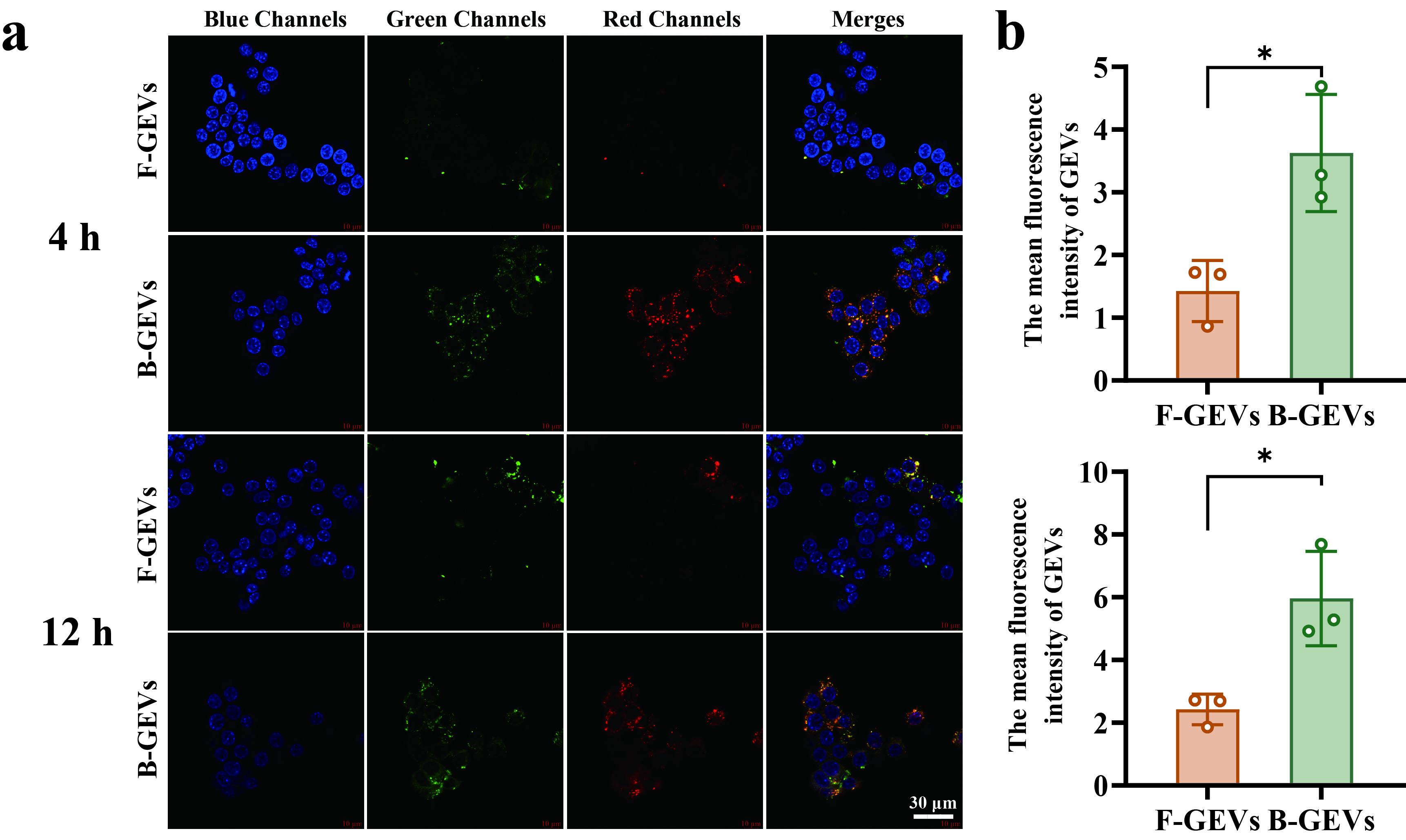

**Figure S3. Uptake of B-GEVs or F-GEVs in RAW264.7 cells detected by confocal microscopy (a-b).** **Blue: nucleus; Green: cell membrane; Red: F-GEVs or B-GEVs.** All data are presented as means ± SD, n = 3. P values were calculated using two-sided one-way ANOVA post-Dunnett’s test; **P* < 0.05.

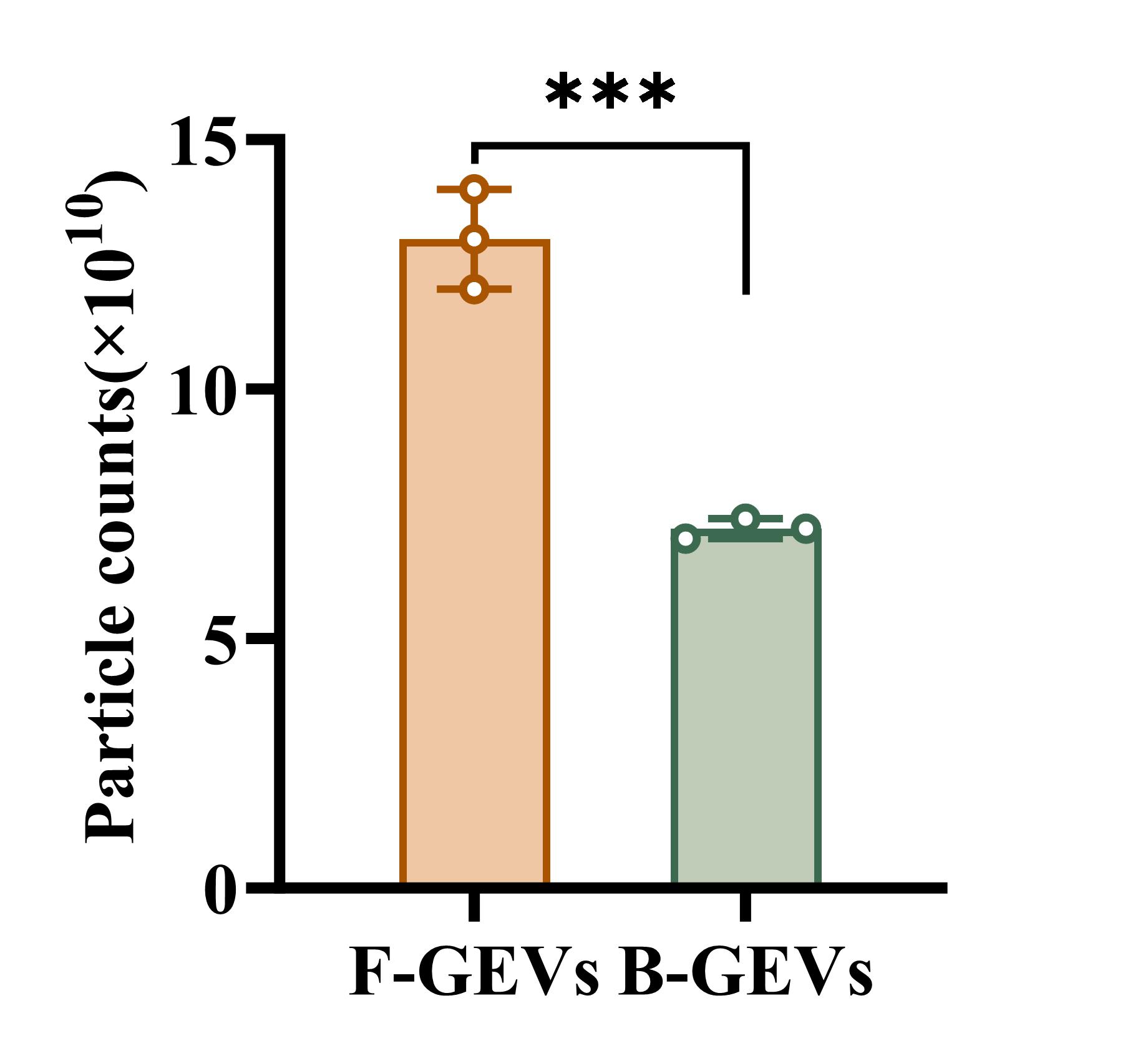

**Figure S4. The particle counts of F-GEVs and B-GEVs determined by NTA.** All data are presented as means ± SD, n = 3. P values were calculated using two-sided one-way ANOVA post-Dunnett’s test; ****P* < 0.001.

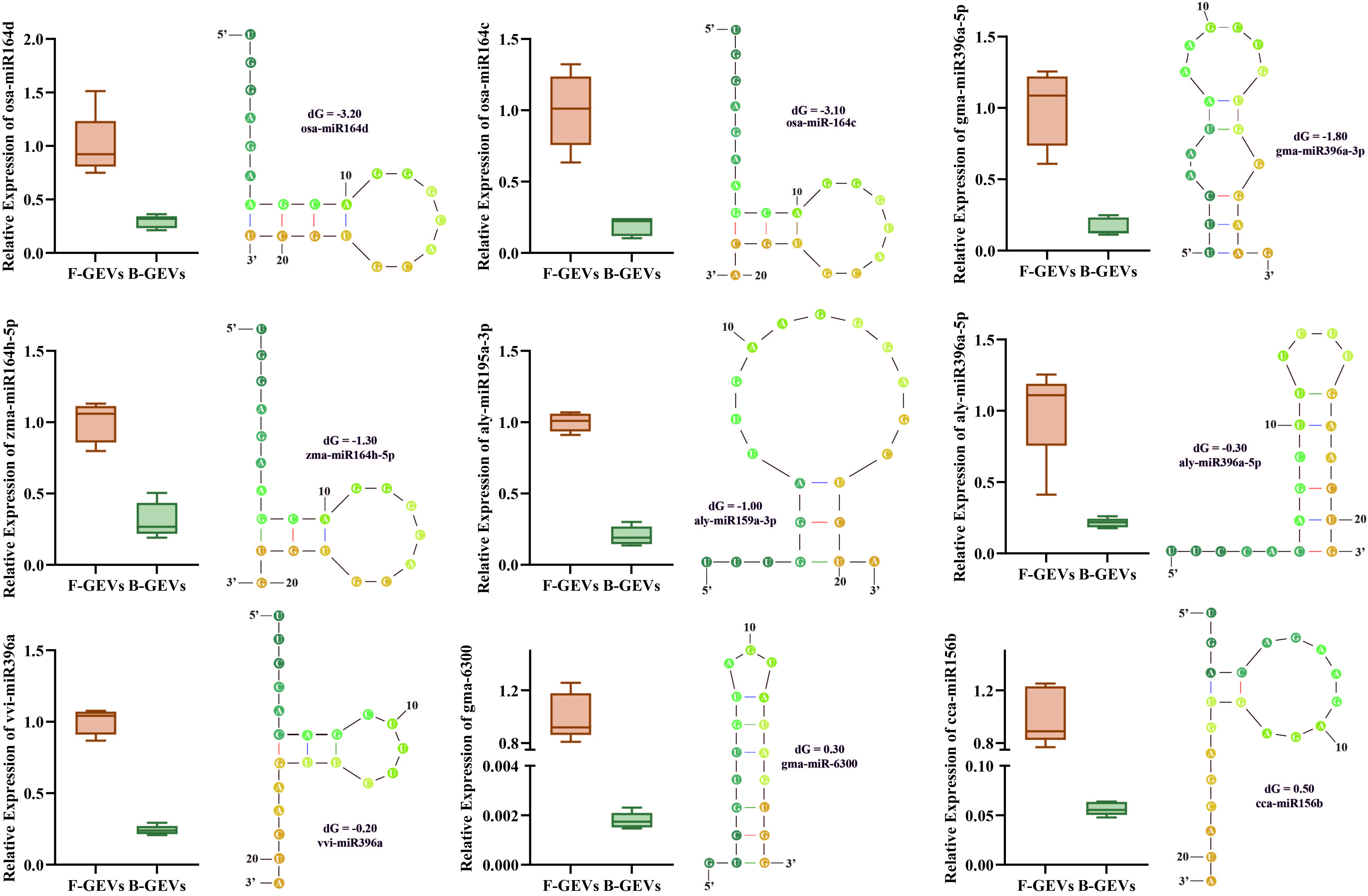

**Figure S5. Relative levels of selected miRNAs found in B-GEVs after boiling.** Relative expression of selected plant miRNAs in F-GEVs and B-GEVs by RT–qPCR (left). Predicted secondary structures with minimum free energy (ΔG) were shown (right). All miRNAs are more abundant in F-GEVs than in B-GEVs.

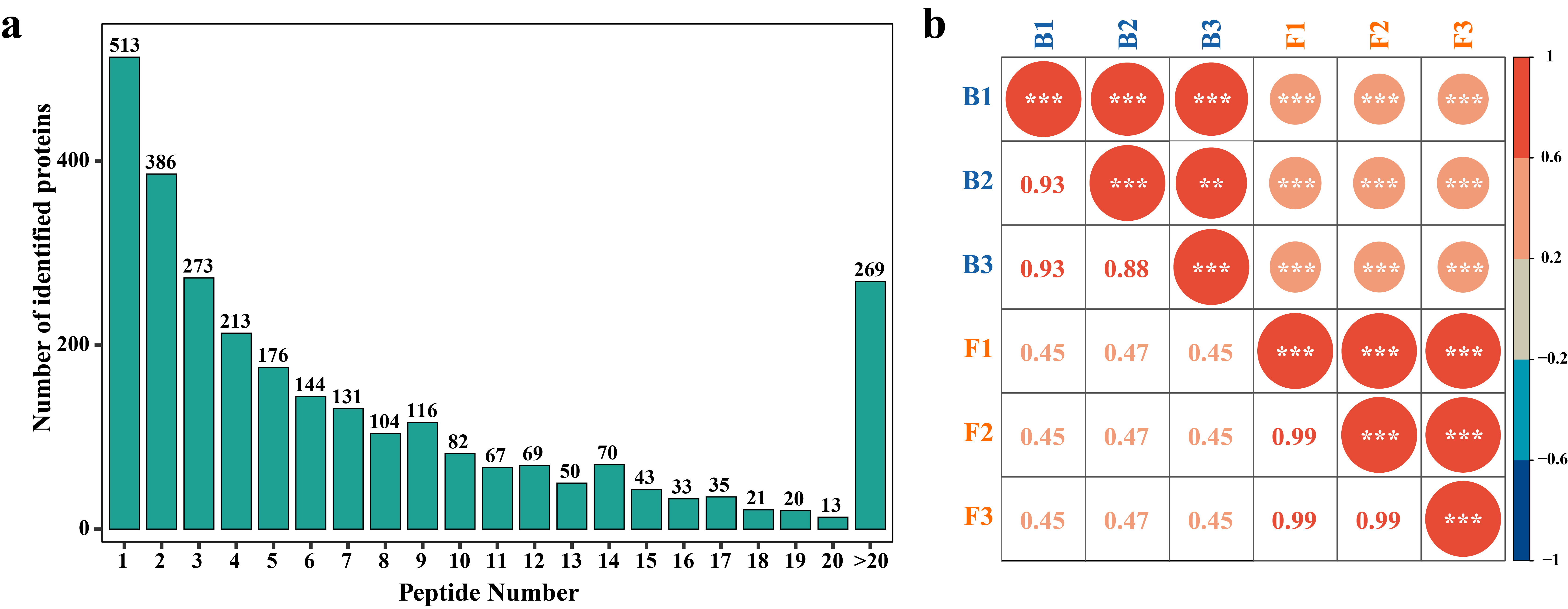

**Figure S6.** **Mass error analysis and correlation analysis of the DAI method based on the LC-MS/MS proteomics workflow**. (**a**), Characterizing the distribution of peptide lengths. (**b)**, Correlation analysis on the different samples for F-GEVs (F1, F2, F3) and B-GEVs (B1, B2, B3).

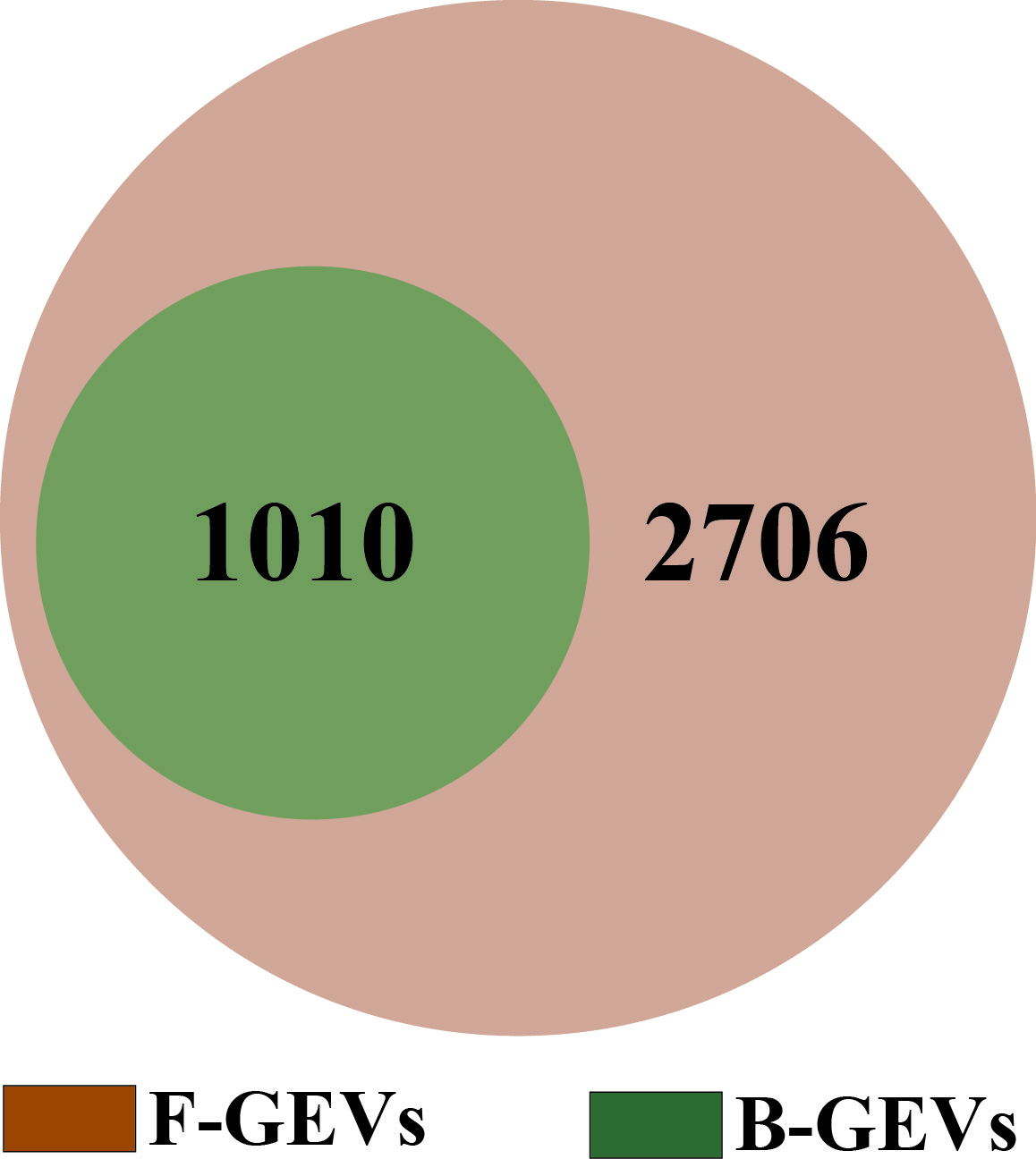

**Figure S7.** **Venn diagram comparing significantly altered proteins as identified.**

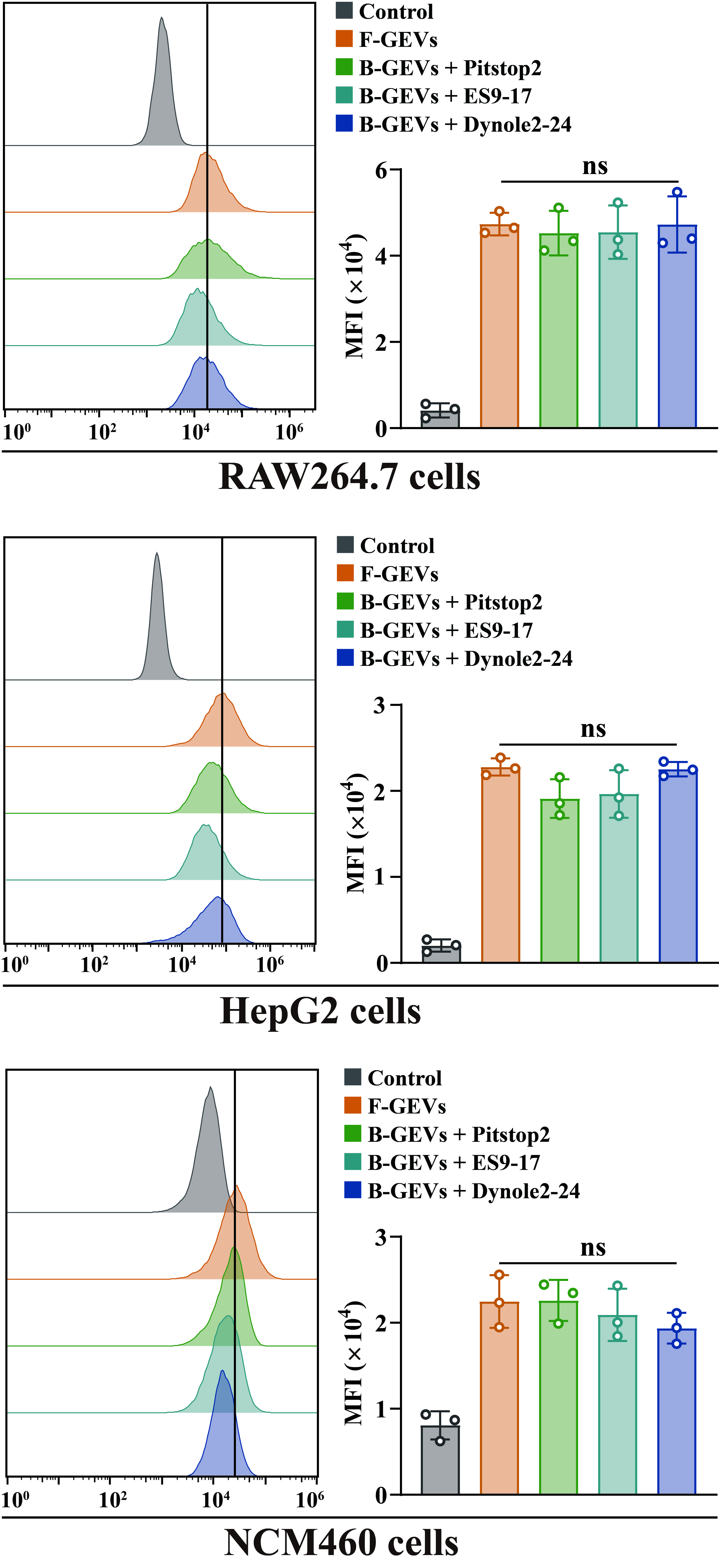

**Figure S8.** **Cellular uptak****e of B-GEVs under different inhibitors of clathrin-mediated endocytosis.** All data are presented as means ± SD, n = 3. P values were calculated using two-sided one-way ANOVA post-Dunnett’s test; ns, non-significant.

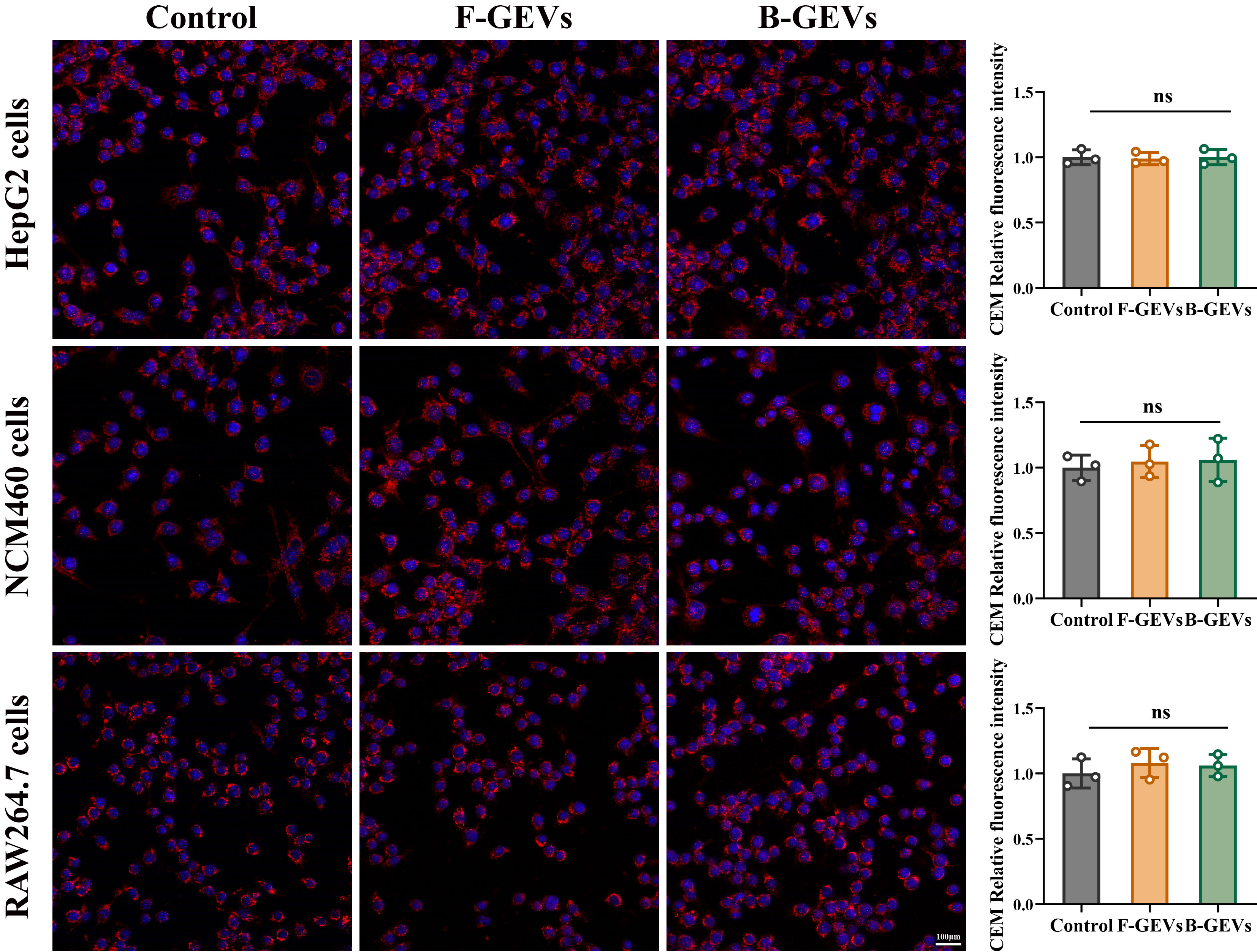

**Figure S9.** **Immunofluorescence analysis of membrane-associated clathrin expression in HepG2 cells, NCM460 cells, and RAW264.7 cells**. Scale bars, 100 μm. All data are presented as means ± SD, n = 3. P values were calculated using two-sided one-way ANOVA post-Dunnett’s test; ns, non-significant.

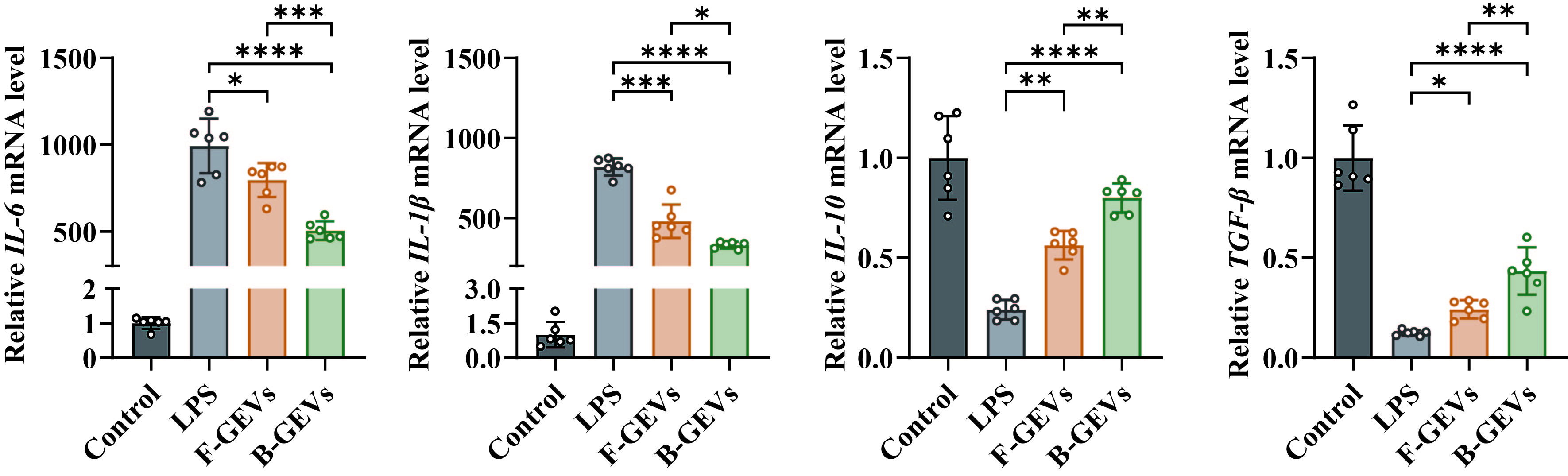

**Figure S10.** **The mRNA expression levels of *IL-6*, *IL-1β*, *TGF-β*, and *IL-10* were measured by RT-qPCR.** All data are presented as means ± SD, n = 6. P values were calculated using two-sided one-way ANOVA post-Dunnett’s test; **P* < 0.05, ***P* < 0.01, ****P* < 0.001, *****P* < 0.0001.

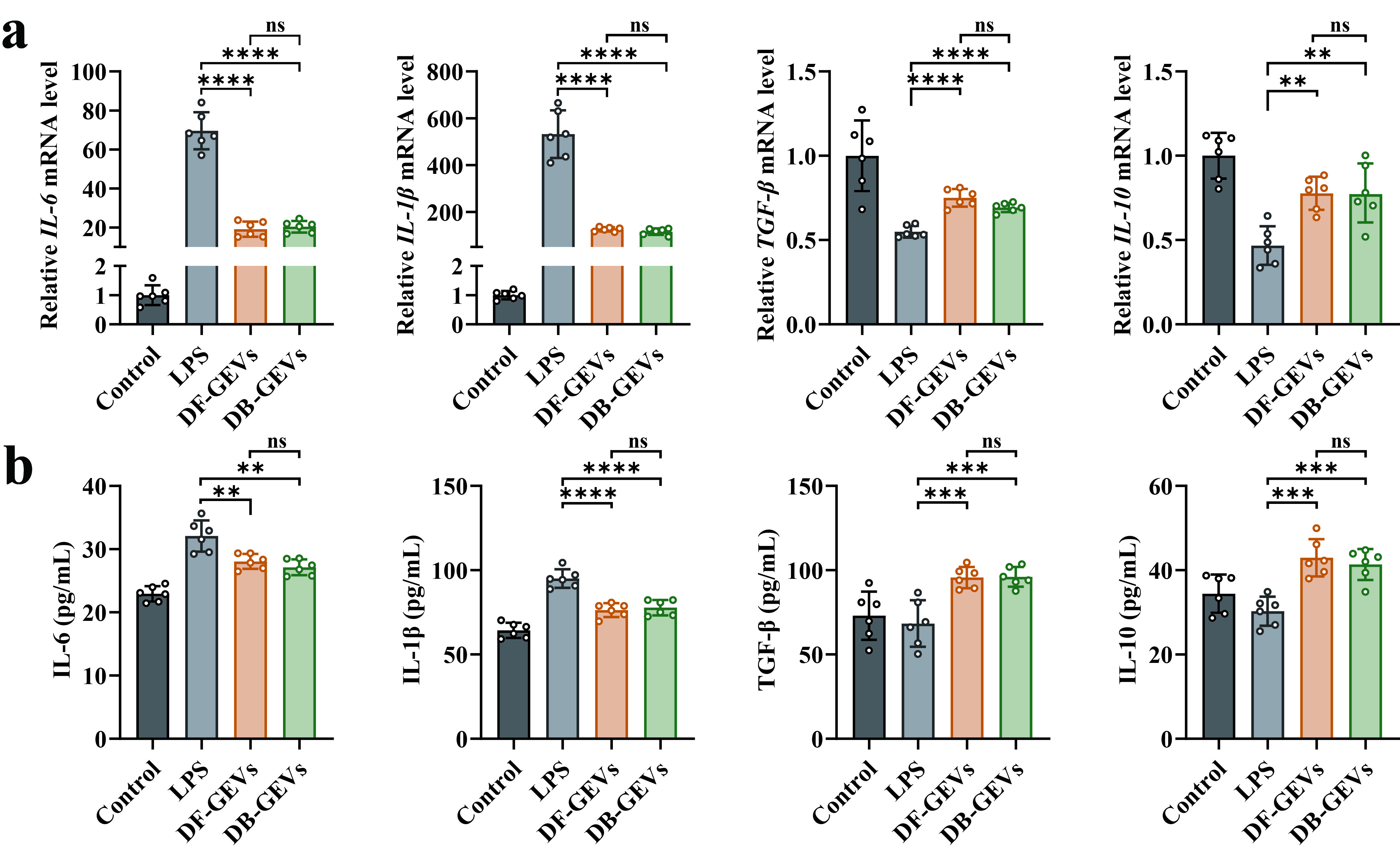

**Figure S11.** **Anti-inflammatory effect of DF-GEVs and DB-GEVs *in vitro***. (**a)**, The mRNA expression levels of *IL-6*, *IL-1β*, *TGF-β*, and *TNF-α* were measured by RT-qPCR (normalized to GAPDH, n=6 biological replicates). (**b)**, ELISA analysis showing the levels of inflammatory cytokines and TGF-β in macrophages transfected with F-GEVs and B-GEVs. All data are presented as means ± SD, n = 6. P values were calculated using two-sided one-way ANOVA post-Dunnett’s test; ***P* < 0.01, ****P* < 0.001, *****P* < 0.0001, ns, non-significant.

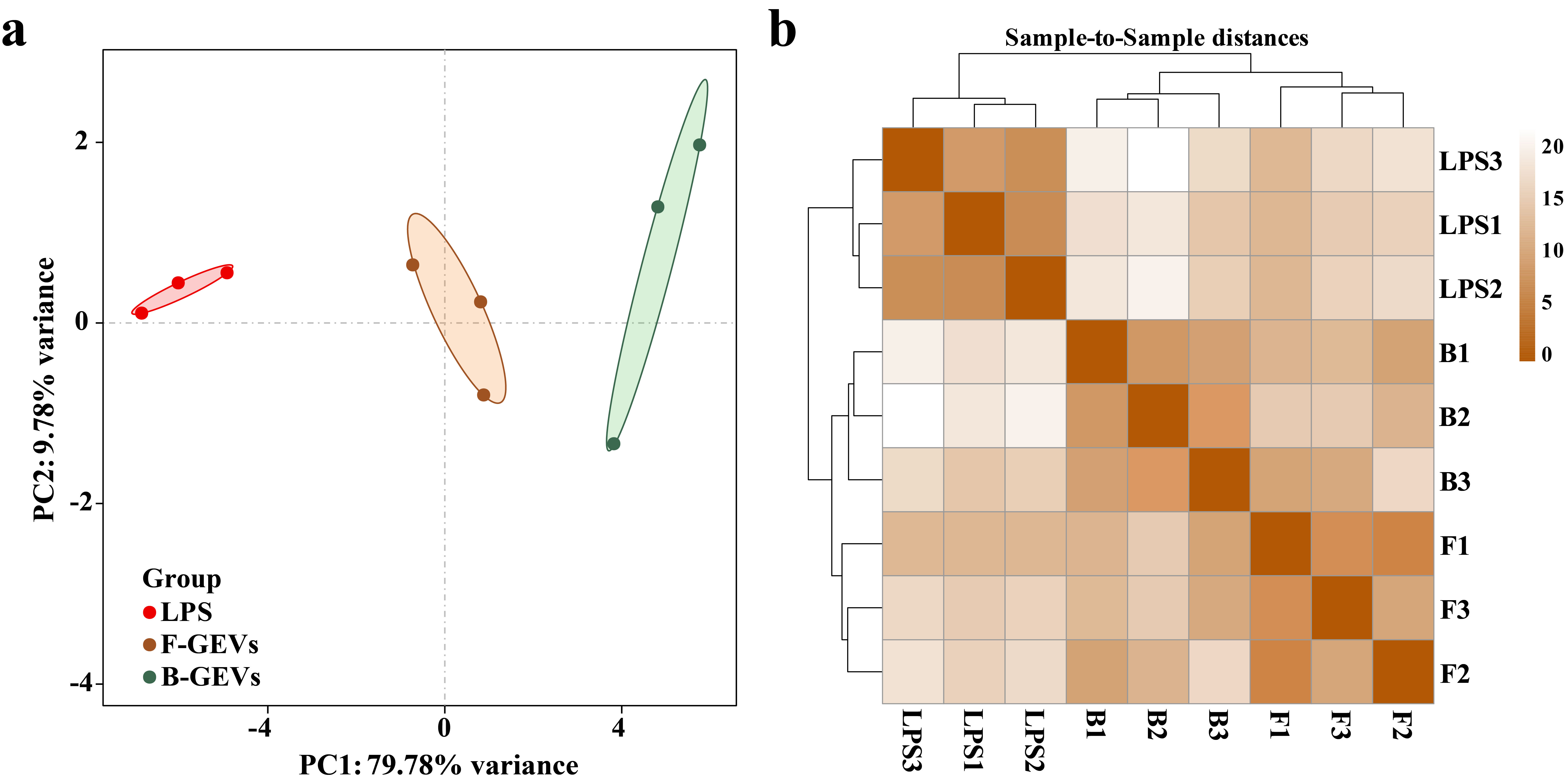

**Figure S12.** **Evaluation of biological replicates and inter-sample correlations.** (**a**), Principal component analysis (PCA) of 9 transcriptomes. (**b**), Hierarchical clustering heatmap illustrating inter-sample relationships.

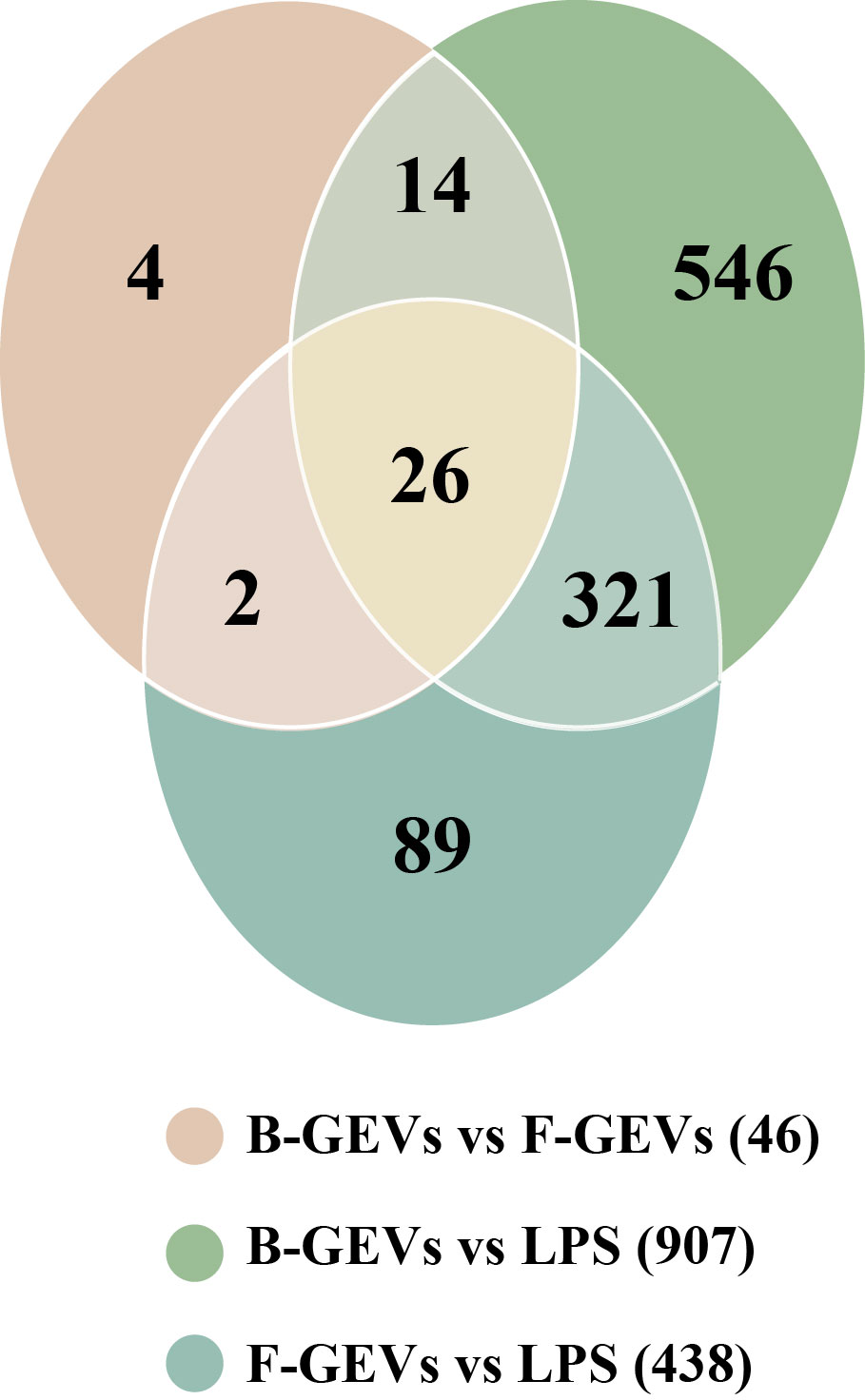

**Figure S13. Venn diagrams visually illustrated the shared and unique differentially expressed genes across the comparison groups.**

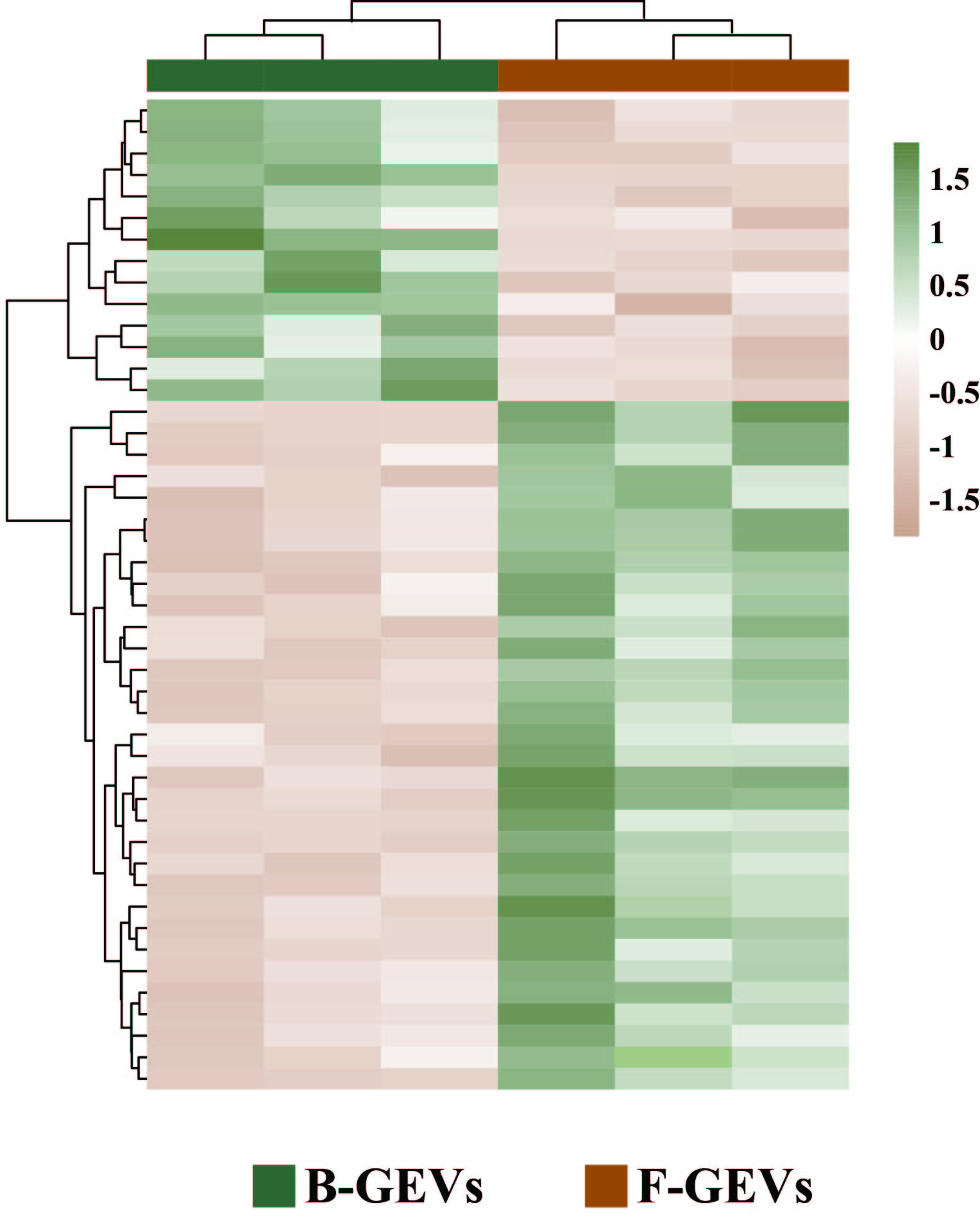

**Figure S14. Heat map of key gene expression showing differentially expressed genes between F-GEVs and B-GEVs.**

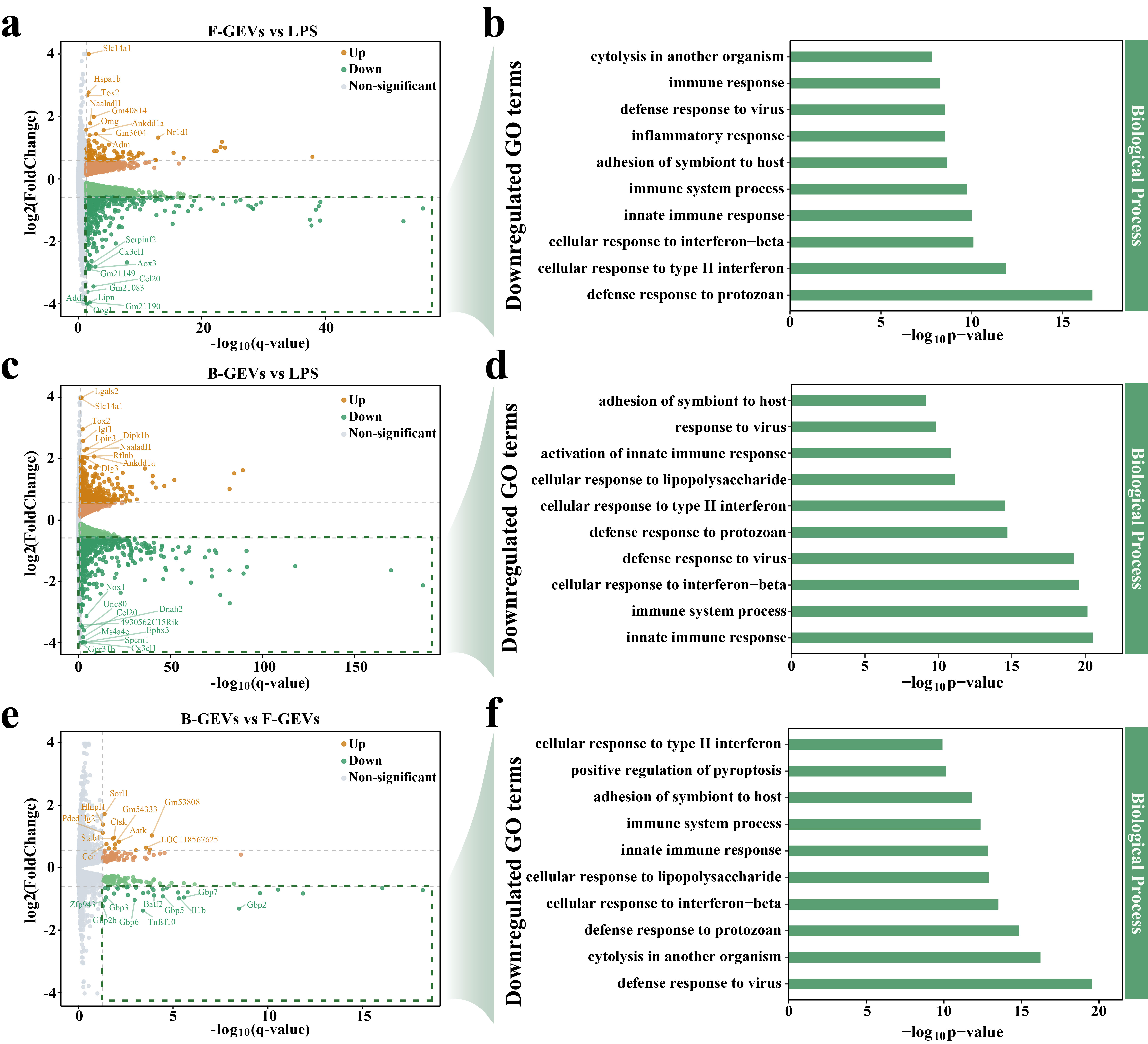

**Figure S15. Downregulation of inflammatory genes in the B-GEVs treatment group.** Comparison of differentially expressed genes between the F-GEVs and LPS group (**a**), the B-GEVs and LPS group (**c**), and the F-GEVs and B-GEVs group (**e**). Top 10 gene GO terms enriched in downregulated genes in (**b**) the F-GEVs vs LPS group, (**d**) the B-GEVs vs LPS group, and (**f**) the F-GEVs vs B-GEVs.

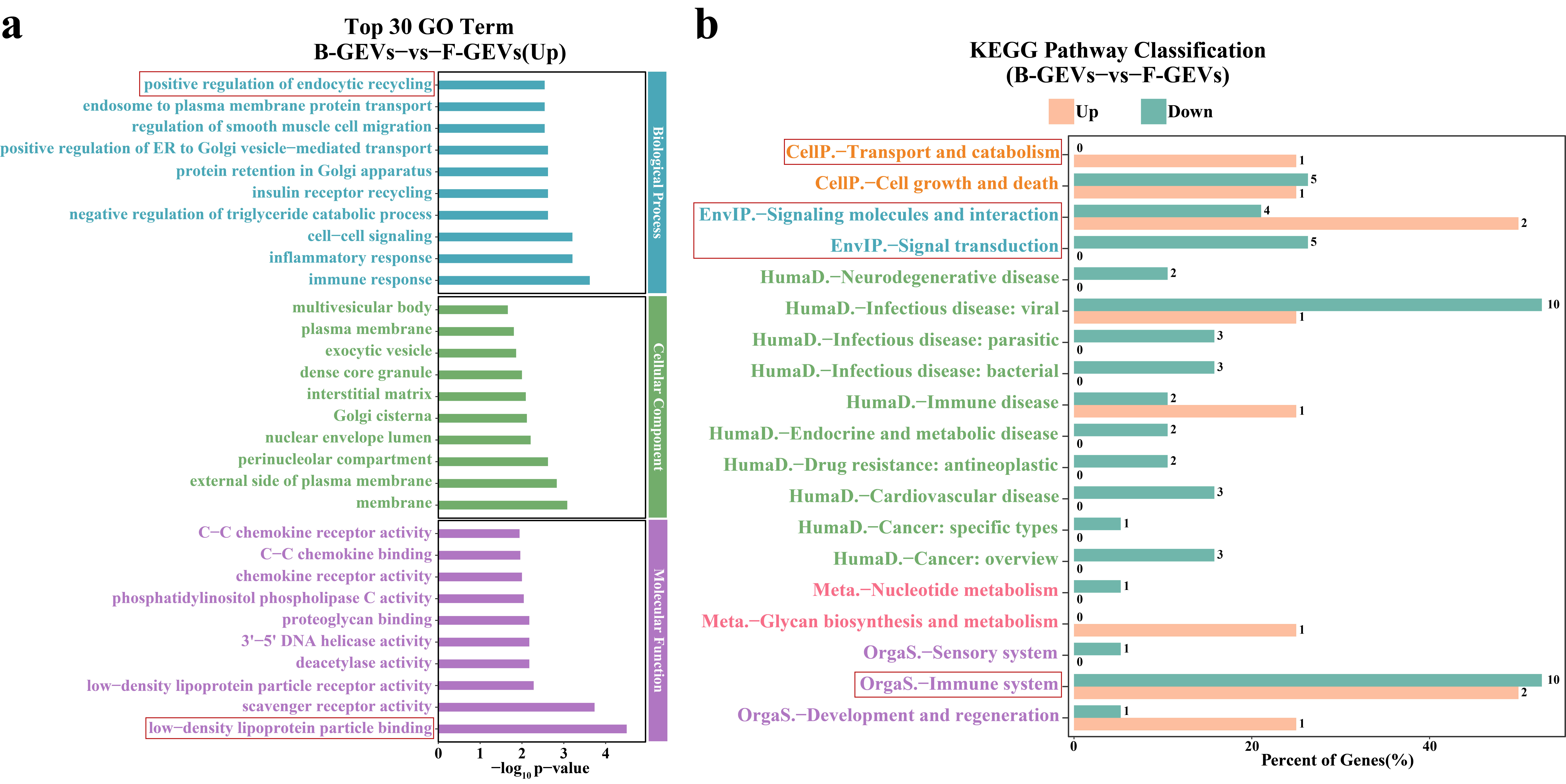

**Figure S16. GO and KEGG pathway enrichment analyses of up-regulated genes in B-GEVs.** (**a**), Top 30 significantly enriched GO terms for genes upregulated in B-GEVs, spanning biological processes (blue), cellular components (green), and molecular functions (purple). (**b**), KEGG pathway classification showing upregulated (orange) and downregulated (green) pathways.

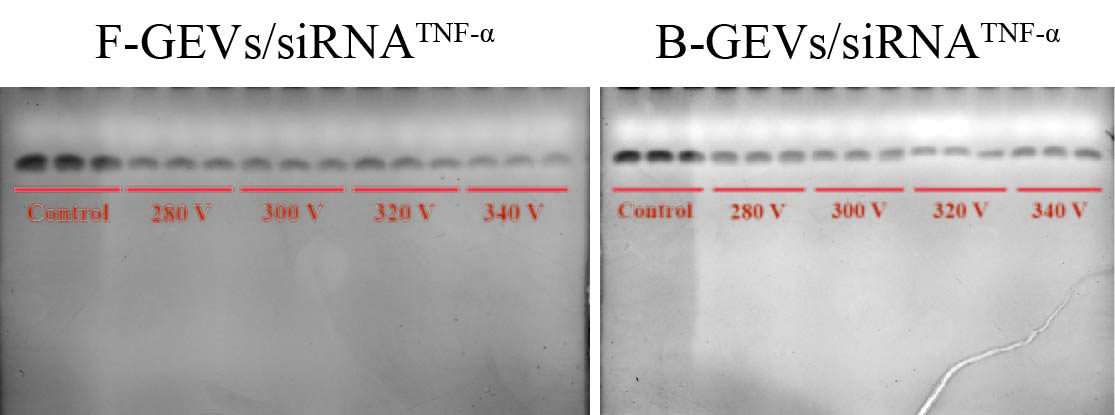

**Figure S17. RNA gel assay for determining siRNA encapsulation efficiency in F-GEVs/siRNA^TNF-α^ and B-GEVs/siRNA^TNF-α^.**

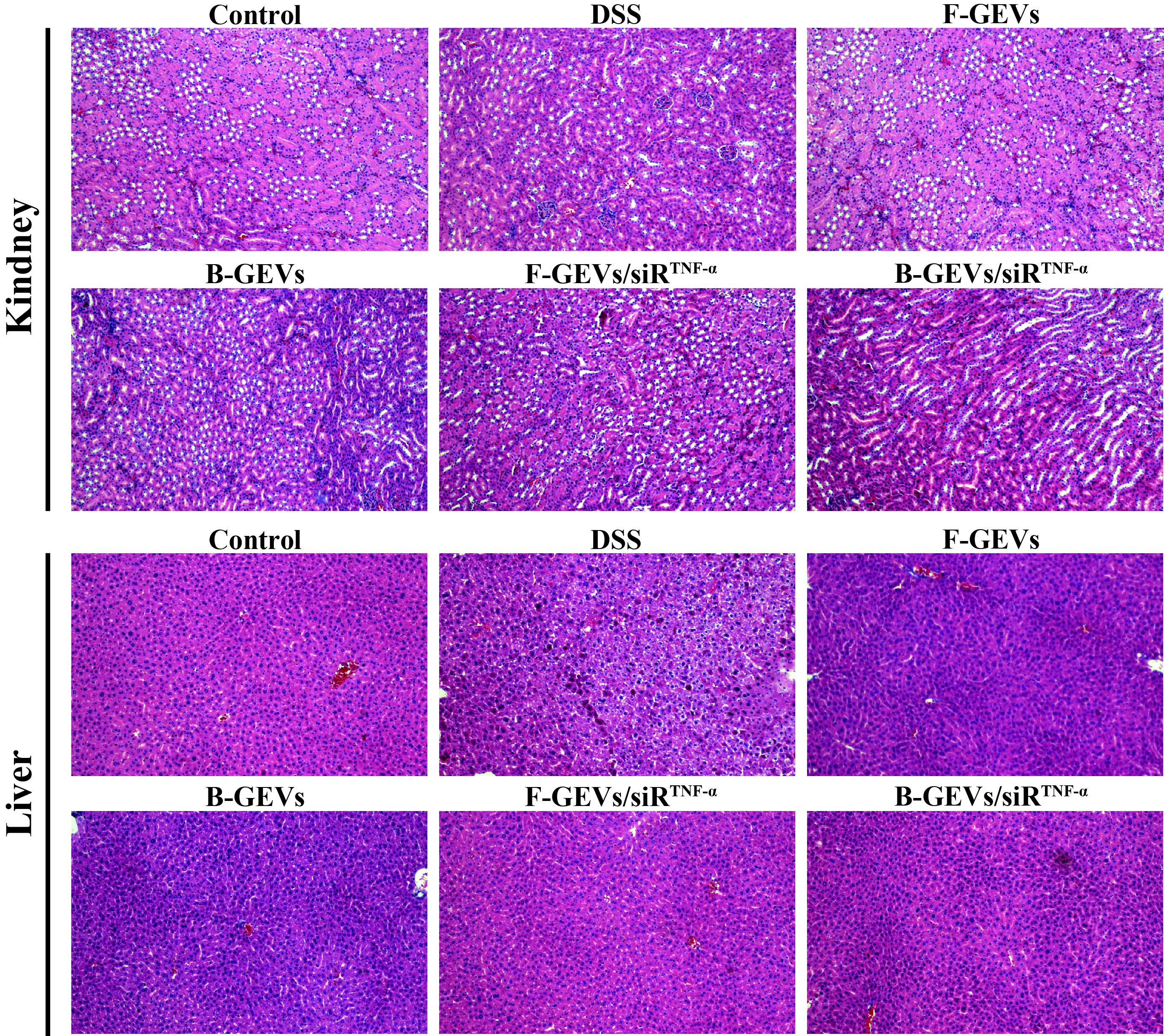

**Figure S18.** **Representative H&E-stained histological sections of kidney and liver tissues.**

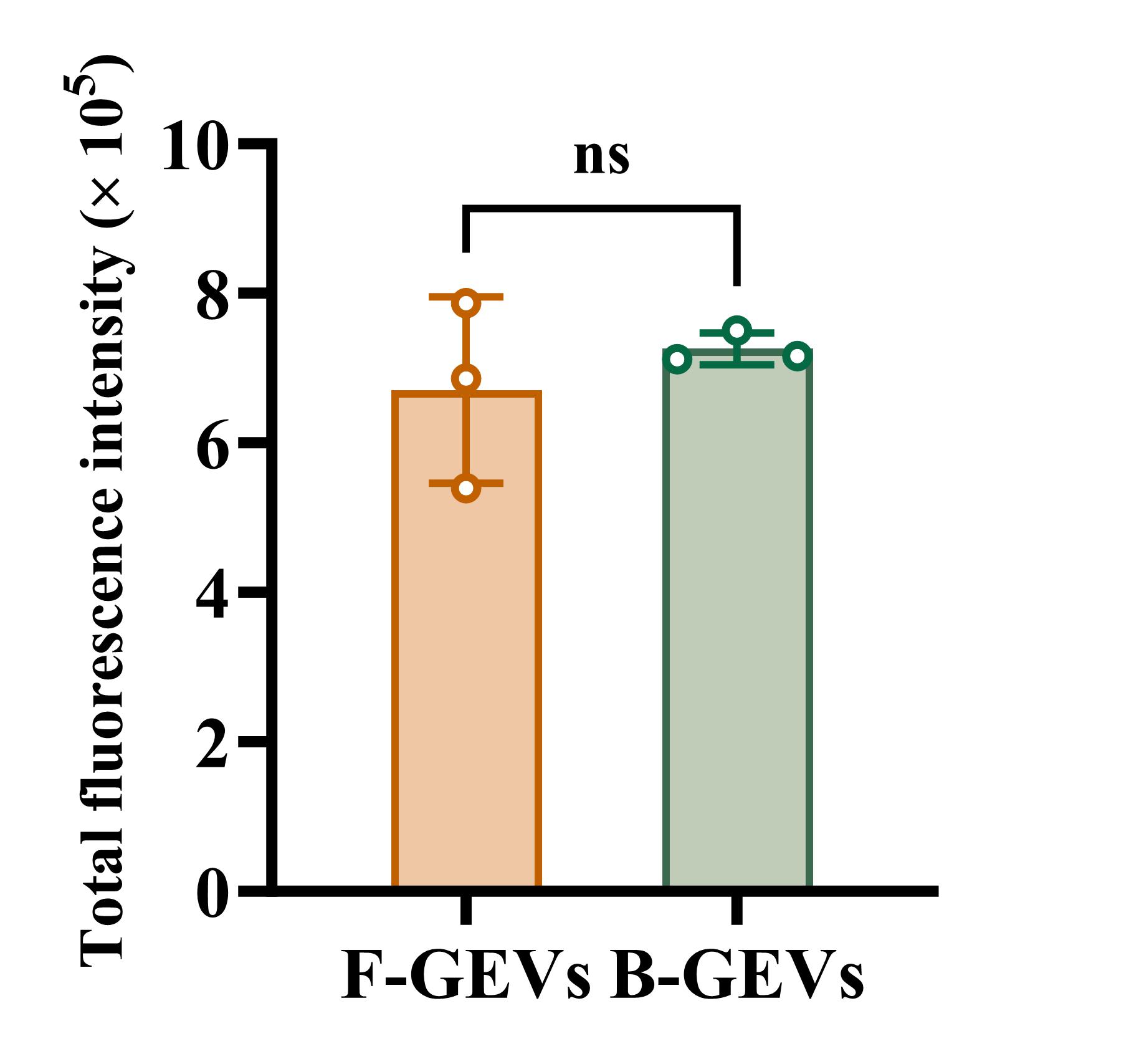

**Figure S19.** **Quantitative analysis of fluorescence intensity from DIO-labeled F-GEVs and B-GEVs.** All data are presented as means ± SD, n = 3. P values were calculated using two-sided one-way ANOVA post-Dunnett’s test; ns, non-significant.

**Table S1. Proteomic profile of significantly up-regulated proteins in B-GEVs compared to F-GEVs.**

| Categories | Accession ID | Number of entries |
| --- | --- | --- |
| Iron Storage Proteins | A0A8J5C078_, A0A8J5GJQ7, A0A8J5C649, A0A8J5C3W0 | 4 |
| Enzymes | A0A8J5H1I9, A0A8J5H5T2, A0A8J5I0Y3, A0A8J5FJR2, A0A8J5GL54, A0A8J5ESJ0, A0A8J5H5L4, A0A8J5HFV4, A0A8J5EW52, A0A8J5KZW9, A0A8J5FEZ1, A0A8J5FZX4, A0A8J5GQ98, A0A8J5GE67, A0A8J5FCX5, A0A8J5F4N3, A0A8J5HY50, A0A8J5F1R5, A0A8J5G393, A0A8J5L7C2, A0A8J5HGC2, A0A8J5F8F4, A0A8J5KEC2, A0A8J5FAP7, A0A8J5G2E0, A0A8J5G723, A0A8J5FWJ6, A0A8J5LKQ6 | 28 |
| Cytoskeleton and structural proteins | A0A8J5HSA3, A0A8J5C5Z1 | 2 |
| Metabolism-related protein | 0A8J5M889, A0A8J5H7F0, A0A8J5HYQ2, A0A8J5G842, A0A8J5G706, A0A8J5HY50, A0A8J5F1R5, A0A8J5G393, A0A8J5L7C2, A0A8J5HGC2, A0A8J5F8F4, A0A8J5KEC2, A0A8J5FAP7, A0A8J5G2E0, A0A8J5G723 | 15 |
| Transporter and membrane proteins | A0A8J5C7J5, A0A8J5BUW5, A0A8J5HMA7, A0A8J5HVS4, A0A8J5CHX9, A0A8J5G7S6, A0A8J5KMM5, A0A8J5I2Z3, A0A8J5EX28, A0A8J5GXD2, A0A8J5HN05, A0A8J5FT37 | 12 |
| Proteases and Inhibitors | A0A8J5LHG8, A0A8J5HXU1, Q5ILG7, A0A8J5GGN5, A0A8J5GYR8, A0A8J5H2Y1, A0A8J5I9T8, A0A8J5EAI2, A0A8J5LFA9 | 9 |
| Signal transduction and regulatory proteins | A0A8J5GZ21, A0A8J5IBG6, A0A8J5GBB3, A0A8J5FLB3, A0A8J5G2E5, A0A8J5G0R2, A0A8J5KRX2, A0A8J5LQN1, A0A8J5GB40, A0A8J5INA7, A0A8J5KL83 | 11 |
| Immunity and defense-related proteins | A0A8J5FY09, A0A8J5KSJ4, A0A8J5FB99, A0A8J5FVG1, A0A8J5L4N4, A0A8J5F2D5, A0A8J5EXW3, A0A8J5GW31, A0A8J5GUY2, A0A8J5KDK3, A0A8J5F1A6, A0A8J5EXT5, A0A8J5EZ66, A0A8J5F5Y1, A0A8J5F663, A0A8J5I0E8, A0A8J5HI52, A0A8J5FVS5, A0A8J5C4N1, A0A8J5G2N9 | 20 |
| Redox and antioxidant proteins | A0A8J5I4A7, A0A8J5KMR2, A0A8J5KU86, A0A8J5GIW4, A0A8J5HCU4, A0A8J5G393, A0A8J5LKQ6, A0A8J5FWJ6 | 8 |
| Molecular chaperones and folding-associated proteins | A0A8J5G7W0, A0A8J5EW23, A0A8J5GKM2, A0A8J5HN05, A0A8J5GFF6, A0A8J5G9K1, A0A8J5C933 | 7 |
| Nucleic acid metabolism-related proteins | A0A8J5BX78, A0A8J5EZD5, A0A8J5FZU0, A0A8J5FS06, A0A8J5C8S5, A0A8J5HQF3, A0A8J5GF55, A0A8J5M7M9, A0A8J5HPS3, A0A8J5ER13 | 10 |
| Multifunctional complex subunit | A0A8J5CEV1, A0A8J5I688, A0A8J5G9K1, A0A8J5FT37, A0A8J5G7S6, A0A8J5CHX9 | 6 |
| Other functionally defined proteins | A0A8J5BUC7 (atpA), A0A8J5BA52 (atpB), A0A8J5GXR6, A0A8J5FBV8, A0A8J5HI46 (CYN), A0A8J5FY97, A0A8J5GZ76, A0A8J5GYZ1, A0A8J5GSD3, A0A8J5BCW2, A0A8J5GG59, A0A8J5BYP6, A0A8J5C5Z8 | 13 |
| Uncharacterized Protein | A0A8J5F812, A0A8J5HCL0, A0A8J5FQM2, A0A8J5LDP7, A0A8J5LNZ5, A0A8J5GIA4, A0A8J5FKI9, A0A8J5F8J5, A0A8J5C6U5, A0A8J5I445, A0A8J5F8H1, A0A8J5LCZ3, A0A8J5L8P8, A0A8J5FZU8, A0A8J5GFF6, A0A8J5L717, A0A8J5HTA2, A0A8J5F3S8, A0A8J5FJR2 | 19 |

Table S2. Sequence of miRNA mimics.

| miRNA | sequence（5’-3’） |
| --- | --- |
| cca-miR156b | UGACAGAAGAGAGUGAGCAUA |
| gma-miR-6300 | GUCGUUGUAGUAUAGUGG |
| osa-miR-164c | UGGAGAAGCAGGGUACGUGCA |
| zma-miR164h-5p | UGGAGAAGCAGGGCACGUGUG |
| gma-miR396a-3p | UUCAAUAAAGCUGUGGGAAG |
| aly-miR396a-5p | UUccacagcUUUcUUgaacUg |
| osa-miR164d | UGGAGAAGCAGGGCACGUGCU |
| aly-miR159a-3p | UUUggaUUgaagggagcUcUa |
| vvi-miR396a | UUCCACAGCUUUCUUGAACUA |

**Table S3. Primer sequence of the GEVs-derived miRNA.**

| miRNA | sequence（5’-3’） |
| --- | --- |
| cca-miR156b | ccgtgacagaagagagtgagcata |
| gma-miR-6300 | cgccgtcgttgtagtatagtgg |
| osa-miR-164c | attggagaagcagggtacgtg |
| zma-miR164h-5p | aatatggagaagcagggcacgtgt |
| gma-miR396a-3p | ccgcttcaataaagctgtgggaag |
| aly-miR396a-5p | cggttccacagctttcttgaactg |
| osa-miR164d | Aatattggagaagcagggcacgt |
| aly-miR159a-3p | ccgtttggattgaagggagctcta |
| vvi-miR396a | CGCTTCCACAGCTTTCTTGAACTA |
| cel-miR-39-3p | TCACCGGGTGTAAATCAGCTTG |

Table S4. Primer sequences for RT-qPCR.

| Gene | primer sequence（5’-3’） |
| --- | --- |
| IL-1β | Forward-GCAGAGCACAAGCCTGTCTTCC  Reverse-ACCTGTCTTGGCCGAGGACTAAG |
| IL-6 | Forward-AATTTCCTCTGGTCTTCTGGAGT  Reverse-GTGACTCCAGCTTATCTCTTGGT |
| IL-10 | Forward-TTCTTTCAAACAAAGGACCAGC  Reverse-GCAACCCAAGTAACCCTTAAAG |
| TGF-β | Forward-GCATTGGCAAAGGTCGGTTT  Reverse-TGCCTCTCGGAACCATGAAC |
| TNF-α | Forward-GCATGATCCGAGATGTGGAACTGG  Reverse-CGCCACGAGCAGGAATGAGAAG |
| NLRP3 | Forward-AAACCCACCAGTGTGCAAGA  Reverse-CAAAGGCCCCTTGTAGCTCA |
| Gbp2 | Forward-GCAAACCCTGGTTCTGCTTG  Reverse-CACATAGTGCAGCTGGTCCA |
| Gbp7 | Forward-AAGGGCATCTGGATGTGGTG  Reverse-ATCTCCTAAGCCCTCCGTGT |
| IL-18 | Forward-ACGGCAAGACCAAGACTCTG  Reverse-GTCACTGCGTTCTCCAGACA |
| GAPDH | Forward-AGGTCGGTGTGAACGGATTTG  Reverse- AGAAGGGTCACTCAGGATAA |
